# Cellular and circuit features distinguish dentate gyrus semilunar granule cells and granule cells activated during contextual memory formation

**DOI:** 10.1101/2024.08.21.608983

**Authors:** Laura Dovek, Mahboubeh Ahmadi, Krista Marrero, Edward Zagha, Vijayalakshmi Santhakumar

## Abstract

The dentate gyrus is critical for spatial memory formation and shows task related activation of cellular ensembles considered as memory engrams. Semilunar granule cells (SGCs), a sparse dentate projection neuron subtype distinct from granule cells (GCs), were recently reported to be enriched among behaviorally activated neurons. However, the mechanisms governing SGC recruitment during memory formation and their role in engram refinement remains unresolved. By examining neurons labeled during contextual memory formation in TRAP2 mice, we empirically tested competing hypotheses for GC and SGC recruitment into memory ensembles. In support of the proposal that more excitable neurons are preferentially recruited into memory ensembles, SGCs showed greater sustained firing than GCs. Additionally, SGCs labeled during memory formation showed less adapting firing than unlabeled SGCs. Our recordings did not reveal glutamatergic connections between behaviorally labeled SGCs and GCs, providing evidence against SGC driven local circuit feedforward excitation in ensemble recruitment. Contrary to a leading hypothesis, there was little evidence for individual SGCs or labeled neuronal ensembles supporting lateral inhibition of unlabeled neurons. Instead, labeled GCs and SGCs received more spontaneous excitatory synaptic inputs than their unlabeled counterparts. Moreover, pairs of GCs and SGCs within labeled neuronal cohorts received more temporally correlated spontaneous excitatory synaptic inputs than labeled-unlabeled neuronal pairs, validating a role for correlated afferent inputs in neuronal ensemble selection. These findings challenge the proposal that SGCs drive dentate GC ensemble refinement, while supporting a role for intrinsic active properties and correlated inputs in preferential SGC recruitment to contextual memory engrams.

**Impact Statement:** Evaluation of semilunar granule cell involvement in dentate gyrus contextual memory processing supports recruitment based on intrinsic and input characteristics while revealing limited contribution to ensemble refinement.

**Major subject area, keywords and organisms:** Semilunar granule cell, inhibition, memory, engram, circuit, hippocampus

## Introduction

The ability of neural circuits to represent unique experiences and events as distinct neuronal representations that can be recalled and updated is fundamental to memory formation. The hippocampal dentate gyrus (DG) is considered central for both novelty detection and to the formation of episodic memories (Hunsaker et al., 2008; Liu et al., 2012; Hainmueller and Bartos, 2020; Danieli et al., 2023). The DG receives dense information from diverse cortical regions through the perforant path projections from the entorhinal cortex (van Groen et al., 2003). Yet relatively few of the numerous closely packed dentate projection neurons, granule cells (GCs), are activated and engage downstream hippocampal circuits. This sparsening of activity is proposed as critical for pattern separation, a process by which the DG helps disambiguate similar memories (McHugh et al., 2007; Hainmueller and Bartos, 2018). Still, the mechanisms that govern how select subsets of neurons are activated during memory formation are not fully understood.

The cellular representations of memories, known as *engrams,* refer to distinct groups of neurons activated during memory acquisition (Semon, 1909; Josselyn et al., 2015; Josselyn and Tonegawa, 2020). Recently, a sparse subset of DG projection neurons, known as semilunar granule cells (SGCs), have been found to be overrepresented among neurons labeled by the expression of the activity dependent immediate early gene (IEG) c-Fos during hippocampus dependent behaviors in TRAP2 reporter mice (Erwin et al., 2020). SGCs, like GCs, have molecular layer dendrites and project axons to CA3 (Williams et al., 2007; Afrasiabi et al., 2022). However, unbiased cluster analyses of morphometric data have revealed that structural features can reliably distinguish SGCs from GCs based on their wider dendritic arbor, greater soma width to length ratio, and more numerous primary dendrites (Williams et al., 2007; Gupta et al., 2020; Afrasiabi et al., 2022). Despite SGCs being estimated to make up only ∼3% of the total GC population (Save et al., 2019), their preferential activation in memory tasks suggests that SGCs may possess unique physiology or connectivity to support recruitment to engrams. However, why SGCs may be preferentially recruited, and whether they shape DG ensemble refinement is unresolved.

There are complementary theories for why certain neurons are selectively activated during memory formation and for how the active cell ensembles may be refined by circuit processes. One hypothesis is that neurons are recruited to memory ensembles based on greater excitability (Yiu et al., 2014; Gouty-Colomer et al., 2016). According to this hypothesis, distinct cohorts of neurons may have higher intrinsic excitability during certain periods; this propensity biases them to fire preferentially in response to inputs and to be recruited into behaviorally activated ensembles (Yiu et al., 2014). Relatedly, it has been suggested that newborn GCs are preferentially recruited to engrams because of their higher excitability (Kee et al., 2007). However, it is not known whether intrinsic physiological features of SGCs, which show sustained afferent driven firing (Larimer and Strowbridge, 2010; Afrasiabi et al., 2022) support their disproportionate representation among behaviorally activated DG ensembles.

In addition to intrinsic properties, neuronal recruitment can be refined by local circuit feedforward or recurrent excitation. One possibility is that glutamatergic interconnectivity aids in engram refinement. Indeed, reports of higher connection probability and strengthening of excitatory synapses between GCs and CA3 pyramidal cells labeled based on IEG expression following fear conditioning (Ryan et al., 2015) support this possibility. Although GCs typically do not innervate other GCs, SGCs have axon collaterals in the molecular layer (Williams et al., 2007; Save et al., 2019), which positions them to potentially form synaptic contacts with GCs. However, whether SGCs directly activate GCs and whether SGCs refine their recruitment to behaviorally active neuronal ensembles remain to be tested. Evaluation of synaptic connectivity between neuronal pairs in ensembles labeled based on IEG expression during memory formation would allow us to test whether recurrent glutamatergic connections support DG ensemble recruitment. Simultaneously, since connectivity between SGCs and GCs is likely to be sparse, this experimental paradigm allows us to address the open question of whether SGCs synaptically activate GCs.

A leading hypothesis for DG circuit refinement of behaviorally active neuronal ensembles, particularly in the context of pattern separation, is through lateral feedback inhibition of surrounding GCs (Walker et al., 2010; Cayco-Gajic and Silver, 2019; Guzman et al., 2021; Borzello et al., 2023). The characteristic robust feedback inhibition in the DG holds promise as a mechanism by which activated engram neurons recruit interneurons to selectively inhibit surrounding neurons (Espinoza et al., 2018). However, this is difficult to reconcile with the exceedingly sparse GC mediated lateral inhibition in recordings from GC pairs (Espinoza et al., 2018; Braganza et al., 2020). It is possible that neurons recruited during contextual memory formation undergo synaptic refinement for better recruitment of lateral inhibition when compared to a naïve circuit. Alternatively, lateral inhibition by neurons in a memory-related ensemble may be largely driven by the recruited SGC populations. SGCs are ideally poised to mediate this effect as their axon collaterals have been shown to form perisomatic synapses on parvalbumin expressing fast spiking basket cells known for their feedback inhibition of GCs (Rovira-Esteban et al., 2020). Consistent with a role for SGCs in supporting feedback inhibition, afferent evoked persistent firing in SGCs is correlated with sustained basket cell and hilar interneuron firing and prolonged inhibitory synaptic barrages in GCs and SGCs (Larimer and Strowbridge, 2010; Afrasiabi et al., 2022). While focal optogenetic activation of a random population of virally labeled GCs elicits robust inhibition in surrounding GCs (Stefanelli et al., 2016), whether the sparse neuronal populations activated during behaviorally driven encoding mediate lateral inhibition of surrounding neurons remains to be tested.

Finally, it is reasonable to posit that precise connectivity of afferent inputs determine downstream activation of a sparse population of DG neurons. Indeed, there is evidence for input-dependent recruitment of neuronal cohorts in the amygdala during fear conditioning (Gouty-Colomer et al., 2016). However, whether shared inputs constrain coactivation of neurons and whether input specificity acts in concert with intrinsic and circuit features to determine which GCs and SGCs are activated is currently unknown. While several studies have focused on DG engram formation (Liu et al., 2012; Ryan et al., 2015), only recently has there been an attempt to explicitly distinguished GCs from SGCs (Erwin et al., 2020). Thus, the specific circuit mechanisms underlying behaviorally relevant DG ensemble refinement during memory encoding and roles of SGCs remain to be determined.

Here we used TRAP2 transgenic mice for c-Fos driven labeling of DG ensembles during behavioral tasks (Guenthner et al., 2013; DeNardo et al., 2019) to label active DG ensembles and undertook ex vivo dual patch clamp and optogenetic recordings in morphologically characterized GCs and SGCs. We use these data to evaluate competing proposals for refinement of cellular ensemble representations in the DG. We specifically focused on differential recruitment of GCs and SGCs in DG ensembles, the potential roles for SGCs in shaping DG circuit processing and refining neuronal ensembles and the role of afferent inputs in shaping DG neuronal ensemble recruitment.

## Results

### Semilunar granule cells are reliably recruited during contextual memory formation

The DG is a primary relay for memory processing (Amaral et al., 2007). However, the mechanisms by which memory-related cellular ensembles are selectively activated during memory encoding are not fully understood. To determine the DG dependent naturalistic behavioral tasks which can recruit a DG ensemble for physiological analysis, we compared the Barnes Maze (BM) and an enriched environment (EE) exposure. We were particularly interested in identifying a behavioral context independent of fear conditioning that activated large cohorts of DG neurons, thereby enabling microcircuit analyses via physiological recordings. Behaviorally activated “*engram*” neurons, referred to henceforth as “labeled neurons”, are neurons in TRAP2 mice induced to express the reporter (tdT or Chr2-YFP) downstream of the activity-dependent IEG c-Fos during BM or EE. “Unlabeled neurons” lack reporter expression. Littermate pairs of TRAP2-tdT mice (4 pairs) were either trained in the BM spatial learning task or exposed to an EE, tasks known to engage the DG. Mice trained in the BM showed progressive decrease in primary latency and primary errors to locate the escape box (Supplemental Fig 1) demonstrating improved performance from acquisition days 1 through 6. Barnes-maze unbiased strategy (BUNS) classification and cognitive scores to assess the use of spatial search strategy (Illouz et al., 2016), revealed that the mice transitioned from using a random or serial search strategy to a spatial strategy as they progressed through acquisition days (Supplemental Fig 1). Both cohorts were induced with tamoxifen during respective behavioral paradigms, on day 6 of BM acquisition or halfway through the 1-day EE exposure, to label active neurons (Fig. 1A-B). Comparison of the number of DG c-Fos expressing (tdT positive) neurons in hippocampal sections from mice one week after tamoxifen induction revealed significantly more tdT labeled neurons following EE exposure than after BM acquisition (Fig. 1C-E; # of tdT labeled cells per slice: EE: 33.90 ± 2.13, BM 13.43 ± 0.90, n=40 slices from 4 animals per group, p=0.0409 by nested t-test). Consistent with previous reports in several other hippocampus-dependent tasks (Erwin et al., 2020), the suprapyramidal (upper) blade of the DG showed more neurons labeled than the infrapyramidal (lower) blade following both BM training and EE exposure (Fig. 1F). To determine whether tagged neurons show task specific reactivation one week after induction, mice were exposed to EE prior to perfusion and sections were immunostained for c-Fos. The distribution of neurons immunolabeled for c-Fos following EE exposure showed no apparent difference between mice previously exposed to BM followed by EE and those exposed to EE twice (Fig. 1Cii, Fig. 1Dii). However, consistent with memory related neuronal tagging, mice with prior exposure to EE showed greater co-labeling of tdT positive neurons with c-Fos immunostaining than mice that were initially trained in the BM task (Fig. 1G; % of co-labeled/total labeled: BM: 2.28 ± 0.46%, EE: 6.8 ± 0.97% p=0.0003 by nested t-test). The results suggest that a cohort of neurons, tagged following EE, reactivate when reintroduced to the same environment, demonstrating memory-specific activation. Therefore, in subsequent experiments, we presumed that cells labeled by task related c-Fos driven reporter expression represent engram cells. Since EE resulted in greater overall DG neuron labeling and stable reactivation of a subset of neurons after one week, we adopted EE as the preferred paradigm to label task-related neuronal ensembles for circuit level analysis.

**Figure 1:**
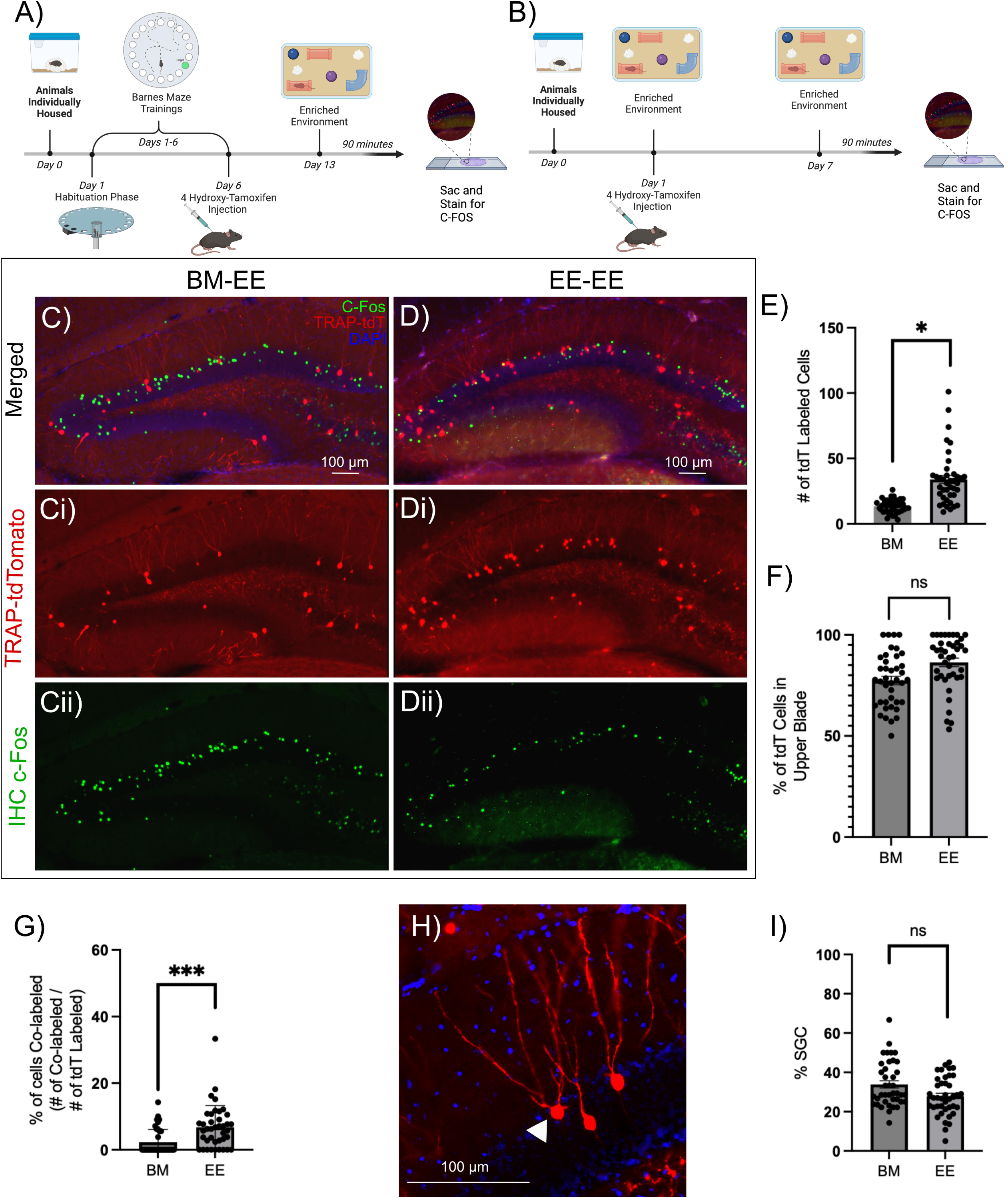
Task associated DG labeled neurons show consistent activation of SGCs and paradigm specific reactivation. A-B) Schematic of experimental timeline for animals trained in the Barnes maze (BM) task followed by exposure to enriched environment (EE), the BM-EE cohort (BM) group (A) and mice housed in EE followed by reintroduction of EE, the EE-EE cohort (EE) group (B). C-D) Representative epifluorescence image of a section from mice one week after induction of tdT labeling (Ci, Di) following BM testing (C) or EE testing (D) and c-Fos immunostaining (Cii, Dii) following subsequent EE exposure. E-F) Quantification of number of tdT labeled cells per slice (E) and summary of proportion of tdT labeled cells in the upper blade of the DG per slice (F). G)Summary of proportion of tdT cells co-labeled with c-Fos (green). H) Representative TRAP-tdT section showing distinct SGC morphology (white arrowhead). I) Plot of % of tdT cells that had morphology consistent with SGCs. * indicates p<0.05, *** indicates p=0.0003 by Nested t-test, n=4 subjects/treatment. Schematics were generated using BioRender under license.

We examined tagged neurons in sections from mice that underwent BM navigation and EE exposure to determine the proportional recruitment of SGCs. SGCs were distinguished from GCs by a trained investigator based on (1) presence of multiple primary dendrites, (2) greater soma width than height, (3) wide dendritic arbor and/or (4) location in or close to the inner molecular layer (Fig. 1H). These criteria were based on our prior studies in which unbiased cluster analysis of GC and SGC morphometric data identified the number of dendrites, soma aspect ratio and dendritic arbor width as the main factors distinguishing the cell types (Gupta et al., 2020; Afrasiabi et al., 2022). The morphology-based classification revealed that 33.86 ± 2.18% of neurons labeled during BM acquisition and 27.83 ± 1.33% during EE exposure were SGCs (Fig. 1I; p=0.1143 by nested t-test, based on 40 sections from 4 mice). Since SGCs represent less than 5% of DG projection neurons (Save et al., 2019), these data suggest preferential activation of SGCs during dentate dependent contextual memory formation. Notably, the proportional recruitment of SGCs labeled following behavior was not different between the BM navigation and EE exposure (Fig. 1I). These findings make a compelling case for leveraging EE exposure to study SGC involvement in dentate-dependent microcircuits.

### Contribution of intrinsic physiology to activity dependent neuronal labeling

To test if the intrinsic physiology of GCs and SGCs labeled during EE differ from their unlabeled counterparts, we performed whole-cell recordings from labeled and unlabeled GCs and SGCs in slices from TRAP2ChR2/eYFP mice one week after tamoxifen induction during EE exposure. Labeled and unlabeled neurons in the granule cell layer and inner molecular layer were visualized under epifluorescence (λ=505 nm) and IR/DIC respectively. Recorded neurons were classified as GC or SGC based on morphology of biocytin filled neurons (Fig. 2A-B) (Williams et al., 2007; Gupta et al., 2020; Afrasiabi et al., 2022). Depolarizing response to blue light activation (0.9mW, λ=470 nm, 10ms) of ChR2 was used to functionally validate cell labeling (Fig. 3E, Fig. 4E). Consistent with earlier studies (Afrasiabi et al., 2022), there was no cell-type specific difference in resting membrane potential (RMP) between GCs and SGCs (Fig. 2C). RMP was also not different between labeled and unlabeled cells within each cell type. Similarly, while SGC input resistance (R_in_) was lower than in GCs, as reported previously (Williams et al., 2007; Afrasiabi et al., 2022), R_in_ of labeled and unlabeled neurons was not different in either cell type (Fig. 2D). Examination of responses to a graded current injection revealed divergence of the firing frequency between GCs and SGCs at current injections >400pA, with GCs showing progressive reduction in frequency with increasing current injection (Fig. 2E-G) due to an apparent depolarization block. Consistently, the firing frequency in response to +520pA current was greater in SGCs than in GCs (Fig. 2H). Again, these cell-type specific differences were maintained in both labeled and unlabeled neurons. The action potential parameters, including threshold, amplitude, half-width, fast afterhyperpolarization (fAHP), medium afterhyperpolarization (mAHP), and latency to first action potential were not different between cell types or labeling of neurons (Supplemental Fig. 2B-G, Supplemental Table 1). Interestingly, GCs showed greater amplitude attenuation during continuous firing, which was not observed in SGCs (Fig. 2I). Once again, these cell-type specific differences were retained in both labeled and unlabeled neurons. Finally, SGCs show higher adaptation ratios (ratio of duration between first two and last two spikes in response to 120 pA current injection), indicating less spike frequency adaptation than in GCs, consistent with previous findings (Williams et al., 2007). Notably, labeled SGCs showed significantly lower adaptation in firing rate than unlabeled SGCs (Fig. 2J; GC_Labeled_: 0.33± 0.075; GC_Unlabeled_: 0.28± 0.056; SGC_Labeled_: 0.78 ±0.076; SGC_Unlabeled_: 0.47 ±0.06; 2-Way RM ANOVA main effect of cell type, p=0.006). In contrast, labeled and unlabeled GCs did not differ in adaptation ratio indicating that the ability to sustain firing may distinguish labeled SGCs. These data support a role for SGC intrinsic physiology, specifically non-attenuating, less adapting, and persistent firing in their preferential labeling during activity dependent neuronal tagging.

**Figure 2:**
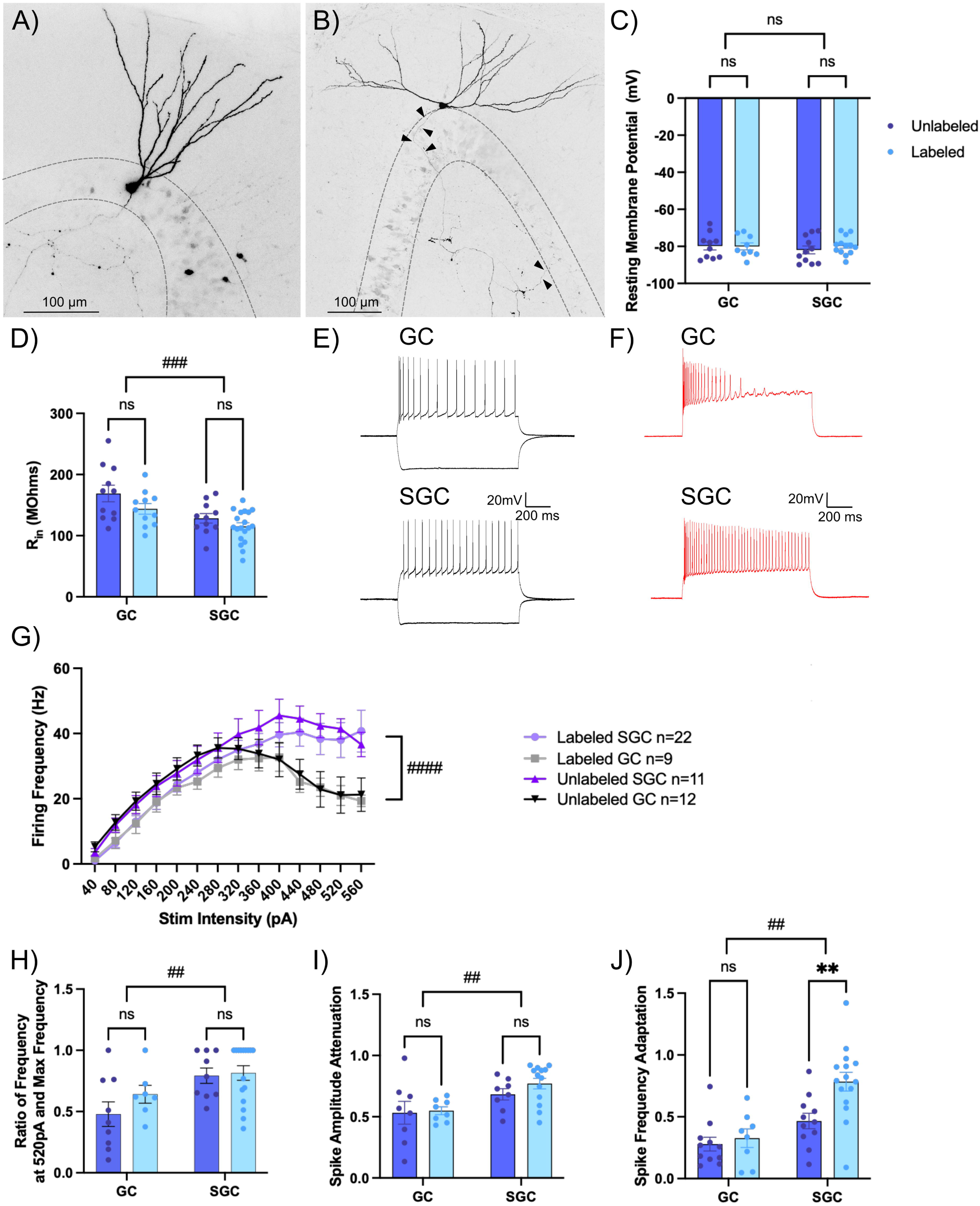
Intrinsic differences in frequency adaptation distinguish labeled SGCs. A-B) Representative images of a biocytin filled GC (A) with a narrow dendritic arbor and a smaller somatic width and a SGC (B) with wide dendritic span, greater somatic width than height, and axonal projections throughout the molecular and granule cell layer (arrowheads). Maximum intensity projections of confocal image stacks are presented as gray scale, inverted images. C-D) Summary plots of resting membrane potential (RMP in C) and input resistance (R_in_ in D) between labeled and unlabeled GCs and SGCs. # indicates p<0.05 for main factor cell type by TW-ANOVA and * indicates p<0.05 for labeled versus unlabeled within cell type by Šídák’s multiple comparisons post hoc test in n=11-19 cells/group. E-F) Representative cell membrane voltage traces in response to +120 and −200pA current injections (E) and +400pA current injection (F) in a GC (top) and SGC (bottom). G) Summary plot of firing frequency in response to increasing current injections in labeled and unlabeled SGCs and GCs. #### indicates p<0.0001 for main factor cell type by 3Way ANOVA n=9-22 cells/group. H-J) Summary plots of firing frequency at 520pA compared to max frequency (H), spike amplitude attenuation calculated as ratio between the amplitude of the 15th spike and 1st spike at a current injection of 400pA (I) and spike frequency adaptation (J). # indicates p<0.05, ## p<0.01 for main factor cell type by TW-ANOVA and ** i ndicates p<0.01 for labeled versus unlabeled within cell type by Šídák’s multiple comparisons post hoc test in n=8-19 cells/group.

**Figure 3:**
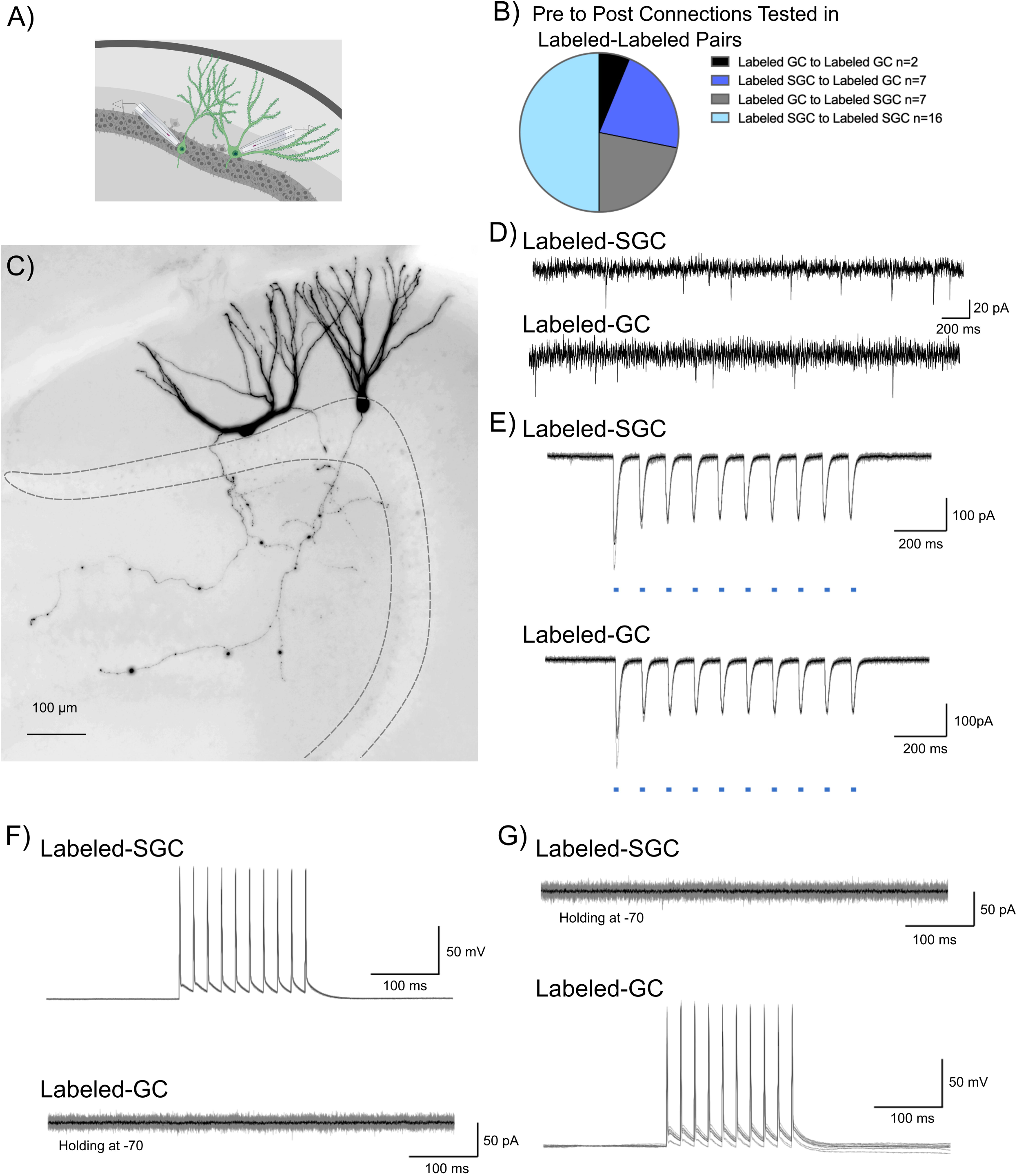
Tagged DG neurons do not support mutual excitatory drive. A) Schematic showing dual patch clamp recording from labeled (green) GC-SGC pair. B) Summary breakdown of cell-type specific connections tested in dual recordings from labeled neurons. C) Representative maximum intensity projection of a confocal image stack of a pair of biocytin filled SGC (left) and GC (right). Images are gray scale and inverted and are overexposed to emphasize the intact axonal arbors in the recorded pair. D) Presence of spontaneous EPSCs in the SGC-GC pair in E-G to verify the presence of excitatory inputs and a healthy circuit. E) Light evoked inward currents validate expression of ChR2 in labeled cell pair. F) Representative traces from a labeled SGC and labeled GC show that depolarization induced firing in SGC (top) failed to evoke EPSCs in a GC (bottom) recorded in voltage clamp. Individual traces are in gray with average trace overlaid in black. G) Depolarization induced firing in GC (bottom) fails to evoke EPSCs in a SGC recorded in voltage clamp (top).

**Figure 4:**
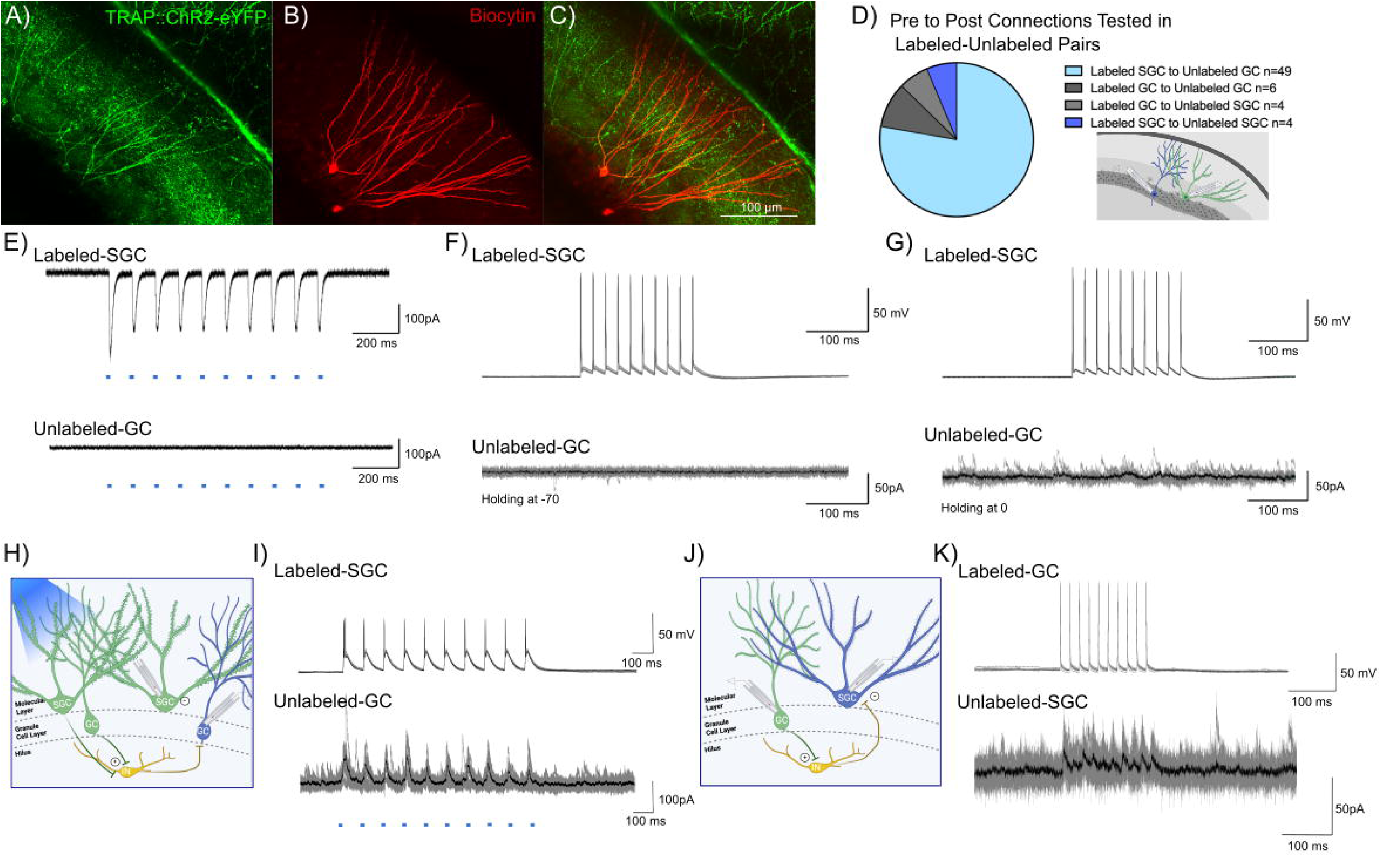
Evidence for DG engram neurons supporting sparse feedback inhibition onto non-engram neurons. A-C) Representative confocal image of eYFP labeled neurons in a TRAP-ChR2-eYFP mouse (A), shows biocytin staining (B) in a pair of recorded Labeled-SGC and Unlabeled-GC. Note co-labeling for eYFP and biocytin in the SGC, while the GC does not colocalize eYFP (C) D) Summary of cell-type specific connections tested in dual recordings from labeled and unlabeled neurons. Inset depicts a schematic showing dual patch clamp recording from a labeled (green) SGC and an unlabeled (blue) GC pair. E) Light evoked currents validate the expression of ChR2 in the Labeled-SGC and lack of response in the Unlabeled-GC. F-G) Representative traces from a Labeled-SGC and an Unlabeled GC show that depolarization induced firing in the Labeled-SGC (top) failed to evoke EPSCs (F) and IPSCs (G) in the Unlabeled-GC. H) Schematic of recording configuration illustrated wide field optical illumination with labeled neurons (green), unlabeled neurons (blue), and local circuit interneuron (yellow). I) Example traces from a recording in which wide field optical stimulation evoked inhibitory responses in the Unlabeled-GC and firing in the Labeled-SGC. Note that the SGC firing by depolarization in the absence of light failed to elicit IPSCs in the same GC. J-K) Schematic with Labeled-GC (green), Unlabeled-SGC (blue), and local circuit interneuron (yellow) (J) and traces from a recorded pair where depolarization of a Labeled-GC elicited inhibitory responses in an Unlabeled-SGC (K).

### Lack of evidence of local feedforward or recurrent excitation between activity driven neuronal ensembles

Unlike GCs, SGCs have axon collaterals in the inner molecular layer and granule cell layer (Fig. 2A,B) (Williams et al., 2007; Save et al., 2019) raising the possibility that they could activate GC dendrites. To test the local feedforward/recurrent excitation hypothesis we conducted dual patch recordings from labeled neuron pairs to identify potential glutamateric interconnections (Fig. 3A,C). Care was taken to ensure that neurons at a depth of 50 µm or more from the surface with visible axons were targeted in order to maximize probability of connections. As noted in Fig. 3B, a majority of the 32 simultaneously recorded neurons examined between labeled neurons were between SGCs (Fig. 3B). However, all other possibilities including GC_Labeled_ to GC_Labeled_, SGC_Labeled_ to GC_Labeled_, and GC_Labeled_ to SGC_Labeled_ were also evaluated. The presence of spontaneous EPSCs in the neurons recorded under voltage clamp served as confirmation of overall circuit and slice health (Fig. 3D). Additionally, optically evoked (0.9 mW, λ=470 nm, 10Hz, 10ms pulses) inward currents provided functional validation of reporter expression (Fig. 3E). Labeled neuronal pairs were tested for glutamatergic synaptic connections by depolarizing one of the neurons in current clamp (400pA, 10 pulses for 10ms at 50Hz) and recording evoked current responses in the other neuron held at −70 mV (Fig. 3F). Recording configuration was reversed to check for connections in both directions (Fig. 3G). Despite the presence of spontaneous EPSCs, none of the 32 labeled neuronal pairs tested, including SGC_Labeled_ to GC_Labeled_ (n=7) and SGC_Labeled_ to SGC_Labeled_ (n=16), showed functional glutamatergic synaptic connections (Fig. 3F-G). Although our data do not eliminate the possibility of direct excitatory connections between labeled neurons, they indicate that glutamatergic interconnections are not critical for activation of DG neuronal ensembles.

### Limited evidence for neurons in activity driven ensembles supporting lateral inhibition

The role of surround inhibition and winner-take-all activation has been proposed as a promising mechanism for establishing memory engrams and mediating dentate processing (Espinoza et al., 2018; Guzman et al., 2021). SGCs, with preferential recruitment in DG engrams, sustained firing, and hilar axon collaterals, are ideally suited to drive robust feedback inhibition (Larimer and Strowbridge, 2010; Walker et al., 2010; Afrasiabi et al., 2022). We examined the possibility that labeled neurons, particularly labeled SGCs, refine GC activity by mediating feedback surround inhibition of unlabeled GCs. We tested this by performing dual recordings from labeled-unlabeled neuronal pairs (Fig. 4A-C). While the majority of our recordings focused on SGC_Labeled_ to GC_Unlabeled_ (49/63 pairs), we also tested for connections between SGC_Labeled_ to SGC_Unlabeled_, GC_Labeled_ to SGC_Unlabeled_ and GC_Labeled_ to GC_Unlabeled_ (Fig. 4D). The ability of wide-field optogenetic activation (0.9 mW, λ=470 nm, 10 pulses, for 10 ms at 10Hz train) to evoke inward currents validated ChR2 expression in labeled neurons. As expected, wide-field blue light stimulation failed to evoke inward currents in unlabeled neurons, confirming the lack of ChR2 expression (Fig. 4E). The recordings also allowed us to test the unlikely possibility that activation of ChR2 expressing labeled neurons evoked synaptic excitation in unlabeled neurons. Wide-field light activation, which likely activates multiple labeled neurons and axon teminals throughout the slice preparation, did not evoke putative polysynaptic EPSCs in unlabeled neurons (n= 63 pairs tested). Consistently, in paired recordings between labeled and unlabeled neurons, current-evoked firing in labeled cells failed to evoke EPSCs in unlabeled cells, (Fig. 4F), underscoring the lack of glutamatergic connectivity between SGCs and GCs.

In paired recordings from labeled SGCs and unlabeled GCs (n=49), depolarization evoked firing in SGCs (400pA, 10ms, 10 pulses, 50Hz) failed to evoke polysynaptic IPSCs in unlabeled GCs (Fig. 4G). Notably, despite the lack of evoked IPSCs, the unlabeled GCs received spontaneous IPSCs indicating that cell and circuit health were not compromised (Fig. 4G). Interestingly, in 1/49 recordings from SGC_Labeled_ to GC_Unlabeled_ pairs, wide-field optogenetic activation at a light intensity that evoked firing in the recorded labeled SGC, evoked IPSCs in the unlabeled GC in the absence of direct synaptic connection between the pairs (Fig. 4H-I). However, in a majority of trials, wide-field optogenetic activation of both labeled GCs and labeled SGCs failed to evoke IPSCs in unlabeled GCs. Activating labeled neurons did not lead to IPSCs in unlabeled neurons in any of the GC_Labeled_ to GC_Unlabeled_ or SGC_Labeled_ to SGC_Unlabeled_ pairs tested. Unexpectedly, we identified one pair (out of 63) in which current induced firing in a labeled GC resulted in robust feedback IPSCs in an unlabeled SGC (Fig. 4J-K). These data identify that labeled GCs can support feedback inhibition of SGCs.

In light of the unexpected paucity in lateral inhibition by engram neurons, we examined whether the circuit connectivity needed to support lateral inhibition is preserved in the slices from mice in which a random cohort GCs were labeled by transfection with AAV5-CAMKIIa-hChR2(H134R)-eYFP to express ChR2 in excitatory neurons. Using a spatial illumination approach we optically activated GC somata in three different decreasing circular regions of interest (ROI) and recorded IPSCs in unlabeled GCs outside the stimulation zone. Focal optical activation of GCs consistently resulted in robust IPSCs in the recorded unlabeled GC with IPSC amplitude decreasing progressively with the size of the ROI (Supplementary Fig 3, p<0.05 by one-way ANOVA, from 5-8 cells from 3 mice). Collectively, although we find that activation of a focal cohort of neurons supports GC lateral inhibition, o data indicate limited evidence for robust lateral inhibition by neurons labeled during EE exposure onto surrounding unlabeled neurons.

### Labeled neuron pairs receive more correlated spontaneous excitatory inputs

Since microcircuit connectivity and intrinsic physiology could not fully account for task related coactivation of neurons, we tested the hypothesis that correlated inputs contribute to ensemble activation. First, we evaluated the contribution of action potential-driven events to GC sEPSCs. Although the frequency of sEPSC in GCs is low, previous studies have identified a substantial contribution of action potential driven events to granule cell sEPSCs in slices from rat (Pernia-Andrade et al., 2012). In GCs recorded under our experimental conditions in mice, the sodium channel blocker TTX (1 µM) consistently increased EPSC interevent interval (Supplemental Fig. 4) by two-fold (2.25±0.3 fold increase in IEI from 5.29±0.84 sec in aCSF to 12.49±2.84 sec in TTX, n=12 cells/3 mice p=0.0005 by paired t-test). These data identify that approximately half the sEPSCs recorded in GCs represent action potential-dependent events and justify analysis of sEPSC in individual neurons and their correlations in neuronal pairs.

To assess whether labeled and unlabeled neurons receive differential glutamatergic drive, we recorded sEPSCs from labeled and unlabeld GCs and SGCs in slices from TRAP2-tdT mice one week after tamoxifen induction following EE (Fig. 5A-B). Recordings from GCs identified that sEPSCs in labeled cells had shorter inter-event intervals (IEI) than unlabeled GCs indicating more frequent sEPSCs (Fig. 5C-D, IEI in s, unlabeled: 5.87 (9.94) n=5 cells from 5 mice; labeled: 2.14 (3.71) n=5 cells from 4 mice, p<0.0001 by K-S test, Cohen’s D = 0.73). Additionally, sEPSC amplitude was also higher in labeled GCs than in unlabeled GCs (Fig. 5D, amplitude in pA,, unlabeled: 16.69 (8.82) n=5 cells from 4 mice; labeled: 20.36 (8.78) n=5 cells from 4 mice, p<0.0001 by K-S test, Cohen’s D = 0.57). As with GCs, labeled SGCs also had lower sEPSC IEI indicating higher frequency (Fig. 5E-F, IEI in s, unlabeled: 2.0 (3.1) n=6 cells from 4 mice; labeled: 1.30 (2.64) n=6 cells from 4 mice, p<0.0001 by K-S test, Cohen’s D = 0.33) and larger sEPSC amplitude (Fig. 5F, amplitude in pA, unlabeled: 15.0 (7.08) n=6 cells from 4 mice; labeled: 19.36 (8.64) n=6 cells from 4 mice, p<0.0001 by K-S test, Cohen’s D = 0.58) than in unlabeled SGCs. Thus, a greater input drive rather than local circuit connectivity distinguishes labeled GCs and SGCs from their unlabeled counterparts.

**Figure 5:**
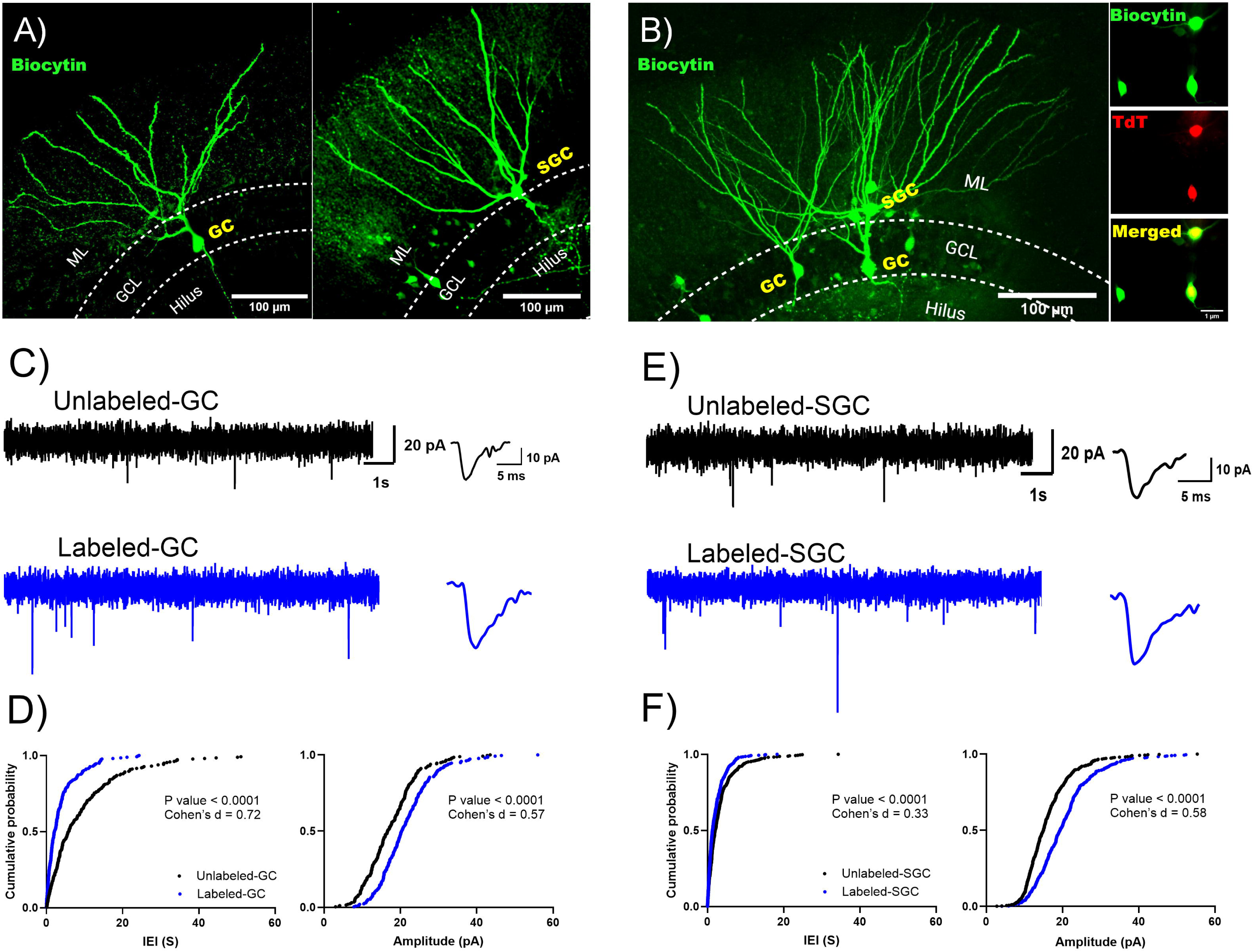
Labeled GCs and SGCs receive more frequent spontaneous excitatory inputs than unlabeled cells. A-B) Representative images of a biocytin filled unlabeled GC (left panel) and SGC (tight panel) (A) and image of a slice in which an unlabeled GC was recorded alongside a labeled GC and SGC (B) Inset in B shows biocytin fill, tdT labeling and merge of the somata to illustrate co-labeling. C) Representative current traces illustrate sEPSCs in an unlabeled (top) and labeled (bottom) GC. Panels to the right: Representative average sEPSCs trace. D) Cumulative probability plot of sEPSC interevent interval (left panel) and amplitude (right panel) in labeled (black) and unlabeled (blue) GC. E) Representative current traces illustrate sEPSCs in an unlabeled (top) and labeled (bottom) SGC. Panels to the right: Representative average sEPSCs trace. F) Cumulative probability plot of sEPSC interevent interval (left panel) and amplitude (right panel) in labeled (black) and unlabeled (blue) SGC. P value by Kolmogorov-Smirnov test is indicated in figure, n=5-6 cells/group. Effect size estimate using Cohen’s D is indicated in the plots.

To evaluate whether temporally correlated inputs contribute to ensemble labeling during EE exposure, we analyzed sEPSC in dual recordings from labeled-labeled (L-L) and labeled-unlabeled (L-U) pairs for temporal correlation of synaptic event times (Fig. 6A-C). Note that since the intent was to determine the input correlation depending on labeling status of the cell pairs rather than based on cell type, we combined datasets for pairs that included GCs and/or SGCs. To assess sEPSC temporal correlation, we defined *peri-occurrence* as the maximum cross correlation of sEPSCs in the two recorded neurons within a *detection window* and *cooccurrence* as the cross correlation of sEPSCs within a more restricted time *bin duration* centered around 0 ms (Fig. 6D). To avoid potential for specious correlations due to differences in event frequency, we sub-selected cell pairs in which the recording durations, event count, and activity rates were not different between L-L and L-U pairs (Supplemental Fig. 5, number of sEPSCs in counts, L-L: 439±52.83, L-U: 412±48.77, p=0.71; recording duration in s, L-L: 498±43.76, L-U: 397±48.0, p=0.15; spike rate in Hz, L-L: 0.89±0.09, L-U: 1.06±0.11, p=0.23).

**Figure 6:**
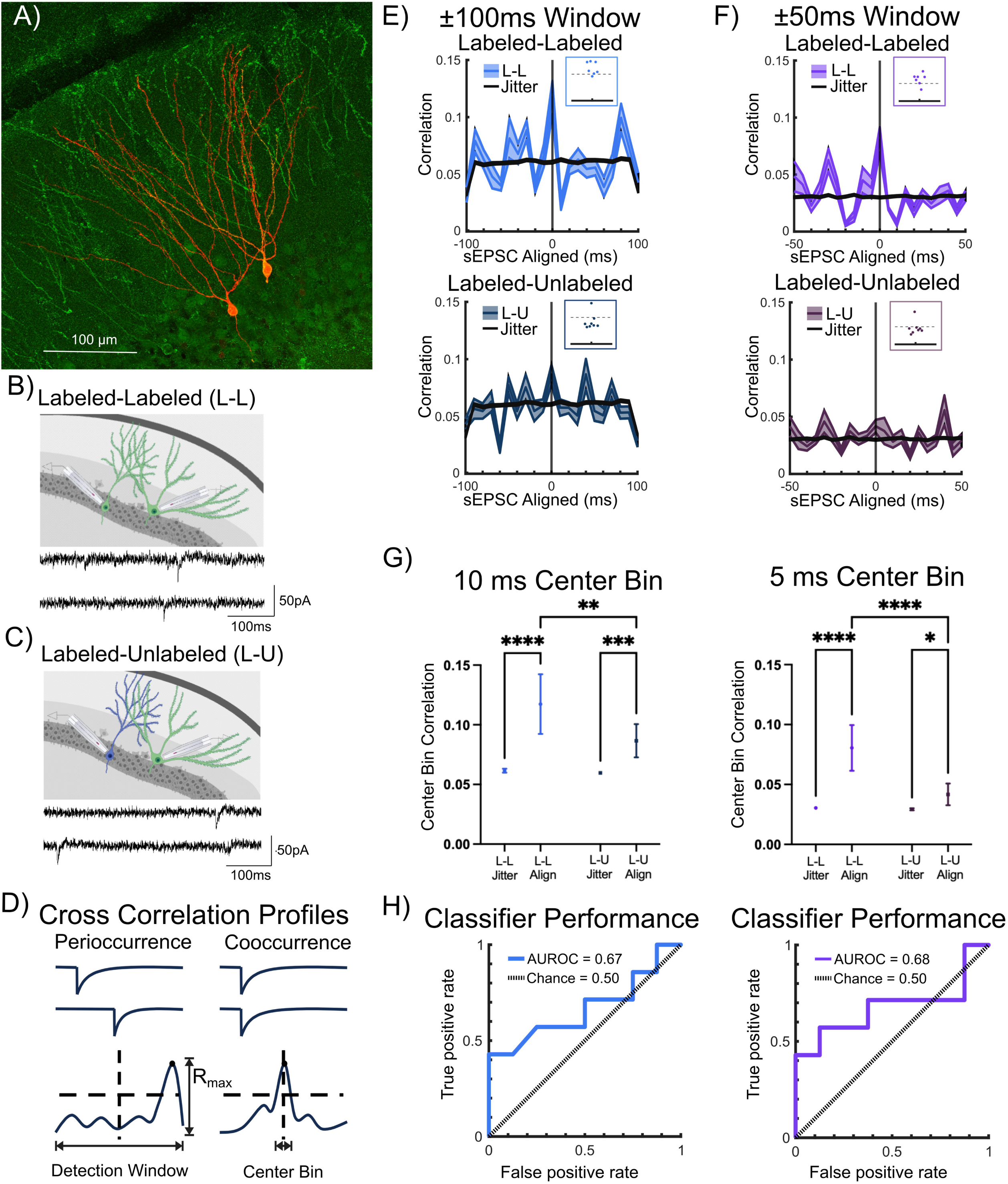
Correlated Spontaneous Excitatory Inputs to Labeled Pairs. A) Representative confocal image of eYFP labeled and biocytin stained neurons in a TRAP-ChR2-eYFP mouse. B) Schematic for Labeled-Labeled (L-L) dual recordings with representative example of spontaneous EPSCs (sEPSCs) in an L-L pair below. C) Schematic for Labeled-Unlabeled (L-U) dual recordings with representative example of sEPSCs in an L-U pair below. D) Schematic for session-wise cross correlation profiles (CCPs) defined by correlations exceeding a 2 standard deviation (SD) threshold above the total mean correlation: EPSC peri-occurrence was tested as event time CCP exceeding threshold within full detection window; cooccurrence was defined as event time CCP exceeding threshold within center bin of detection window. E) CCP from recordings from L-L pairs analyzed with ±100 ms detection window (bright blue, n=7). Overlaid jittered data (black) was developed by appending the event timing of one cell with a randomized lead/lag of +/- 0.5 s for 100 iterations (Top panel). Inset: Plot of maximum correlations (Rmax) in relation to the dashed line representing 2xSD = 0.15. CCP in recordings from L-U pairs analyzed with ±100 ms detection window (dark blue, n=8). Corresponding jittered data, developed as detailed above, is overlaid in black (Bottom panel). Inset: Plot of Rmax in relation to the dashed line representing 2xSD = 0.15. F) CCP from sessions with recordings from L-L pairs analyzed with ±50 ms detection window from L-L pairs (bright purple, n=7) with jittered data developed as detailed above is overlaid in black (Top panel). Inset: Plot of Rmax in relation to the dashed line representing 2xSD = 0.10). CCP from recordings in L-U pairs analyzed with ±50 ms detection window (dark purple, n=8) with corresponding jittered data overlaid in black (Bottom panel). Inset: Rmax in relation to the dashed line representing 2xSD = 0.10. G) Comparison of center bin correlation between L-L versus L-U pairs in aligned (Align-recorded) versus jittered (Jitter-simulated) data, analyzed using ±100 ms detection window (left, colors as in E) and using ±50 ms detection window (right, colors as in F). H) Center bin classifier performance (solid line) compared to chance performance (dashed line, colors as in E and F respectively) plotted as area under the receiver-operating characteristic (ROC) curve (AUROC) between L-L (true positive rate) and L-U (false positive rate) for analysis using ±100 ms detection window (left panel) and for analysis using ±50 ms detection window (right panel). Data presented as mean±SEM (dual recording sessions), * indicate p<0.05; ** indicates p<0.01 *** indicates p<0.001, **** indicates p<0.0001; TW-ANOVA with Šídák’s multiple comparisons post hoc tests.

We selected detection windows of ±100 ms and ±50 ms with 10 ms and 5 ms bin width, respectively, to develop cross correlation profiles (CCPs) of L-L and L-U sEPSC event times as detailed in the methods. Example recording sessions with CCP for cooccurrence in L-L pairs, no coincidence in L-U pairs, and peri-occurrence in L-L pairs, analyzed using a ±100 ms detection window are illustrated in Supplemental Fig. 6). A predetermined threshold of 2 standard deviations (2SD) above the mean correlation was adopted to assess potential differences in temporal correlation between L-L and L-U pairs. Peri-occurrence, quantified as the maximum cross correlation in the detection window (R_max_), was significantly higher in L-L than in L-U pairs for the ±100 ms detection window (Fig. 6, R_max_ in ±100 ms window, L-L: 0.184±0.014, L-U: 0.126±0.016, p=0.018 by t-test in 7 L-L and 8 L-U pairs). While peri-occurrence in the ±50 ms detection window trended higher in L-L pairs, this was not significant (Fig. 6, R_max_ in ±50 ms window, L-L: 0.124±0.013, L-U: 0.093±0.014, p=0.12 by t-test in 7 L-L and 8 L-U pairs). Notably, the R_max_ in sEPSC event times from 6/7 L-L pairs crossed the threshold within each detection window and failed to do so in 7/8 recordings in L-U pairs. These findings were consistent regardless of whether we adopted ±100 ms or ±50 ms detection windows (Fig. 6E-F insets).

To determine if event correlations in neuronal pairs deviated from random, event time correlations in the recorded (*temporally aligned*) data were compared with the correlations developed from corresponding temporally *jittered* data sets (Fig. 6E-F). Event time center bin correlations in the recorded (*temporally aligned*) data sets were significantly higher than in the *jittered* data sets for both detection windows (Fig. 6G main effect of data alignment, 10 ms bin: F(1,211)=63.16, p<0.0001 by TW-ANOVA, 5 ms bin, F(1,211)=71.60, p<0.0001 by TW-ANOVA). Cooccurrence, quantified as the correlation in the center bin, was higher in L-L pairs both within the 10 ms bin (Fig. 6E,G, 10 ms center bin correlation, L-L: 0.117±0.025, L-U: 0.087±0.014, p=0.0075 by TW-ANOVA in 7 L-L and 8 L-U pairs) and within the 5 ms bin (Fig. 6F, G, 5 ms center bin correlation, L-L: 0.081±0.019, L-U: 0.042±0.009, p<0.0001 by TW-ANOVA in 7 L-L and 8 L-U pairs

Finally, we evaluated the ability of cooccurring events to predict an L-L versus L-U recording session. The receiver operating characteristic (ROC) curve of true versus false positive rates defined the area under the ROC curve (AUROC) for the center 10 ms and 5 ms bin across the ±100 ms and ±50 ms detection windows, respectively (Fig. 6H). Center bin classification performed better than chance at predicting whether a recorded session was L-L versus L-U (AUROC_Chance_ = 50%; AUROC_10msCenter_ = 66.96%; AUROC_5msCenter_ = 71.43%). Thus, the 10 ms and 5 ms center event correlations were predictive of whether a pair of recorded neurons were likely to be a pair of labeled neurons or a labeled-unlabeled pair. Together these results support a role for correlated inputs in driving shared neuronal activation during contextual memory formation.

## Discussion

The recent characterization of semilunar granule cells as a unique dentate projection neuron subtype overrepresented amongst behaviorally recruited DG neurons has raised the intriguing possibility that SGCs may play a distinct role in shaping DG ensemble activity (Walker et al., 2010; Erwin et al., 2020; Afrasiabi et al., 2022). Here we evaluated competing hypotheses involving mechanisms governing recruitment of GC and SGC populations during a behavioral experience. Our data identify that *intrinsic properties of SGCs,* specifically their *less adapting firing* characteristics, likely enable preferential recruitment of SGCs among neurons labeled based on IEG expression. At the circuit level, neurons activated during a behavioral experience received more frequent and larger excitatory synaptic input than those not engaged in the task. Notably, neurons in a shared DG ensemble receive *more correlated spontaneous excitatory inputs* than neurons without shared activation, suggesting a role for common afferent inputs in behaviorally driven ensemble recruitment. Whereas GCs and downstream CA3 neurons with shared behavioral activation develop preferential glutamatergic connectivity (Ryan et al., 2015), we found no evidence for local feedforward or recurrent excitation in DG neurons labeled as part of a memory trace. Unexpectedly, although lateral inhibition has been proposed as a mechanism for dentate engram refinement (Stefanelli et al., 2016), our experiments revealed that activation of labeled DG neurons rarely drove inhibitory synaptic currents in unlabeled neurons. Interestingly, we find approximately a third of the projection neurons activated as a part of a dentate dependent spatial navigation task and during exposure to enriched environment had morphology consistent with SGCs. It is possible that the reduced action potential accommodation and attenuation in SGCs contributes to enhanced SGC firing and c-Fos expression during afferent activation. This greater activity dependent c-Fos expression in SGCs may result in their preferential labeling as part of neuronal ensembles activated during behavioral tasks. Together, these results identify that behaviorally relevant activation of SGCs and GCs in DG neuronal ensembles are determined by a combination of sustained firing characteristics of SGCs, enhanced glutamatergic inputs and shared afferent drive rather than by selective circuit level refinement by recurrent excitation or by lateral inhibition.

Studies evaluating mechanisms of ensemble recruitment during fear and aversive memory encoding, by experimentally enhancing excitability or CREB expression in a sparse population of amygdala neurons, have proposed that neurons with higher excitability outcompete neighboring cells for allocation to behaviorally activated neuronal ensembles (Han et al., 2007; Zhou et al., 2009; Sano et al., 2014; Yiu et al., 2014; Gouty-Colomer et al., 2016). However, our data revealed no difference in intrinsic physiology and active properties between labeled and unlabeled GCs. This is consistent with a previous report that firing threshold and input resistance of GCs labeled during fear conditioning and recorded 24 hours later were not different from unlabeled GCs (Ryan et al., 2015), although GCs and SGCs were not explicitly distinguished. In addition to previously reported less adapting firing in SGCs than in GCs (Walker et al., 2010; Afrasiabi et al., 2022), we find reduced spike amplitude attenuation in SGCs resulting in more sustained firing, particularly in response to large current injections. Moreover, in the first direct comparison of behaviorally recruited and unlabeled SGC, we identify that labeled SGCs had lower spike frequency adaptation than unlabeled SGCs, indicating that sustained firing may predispose SGCs to activation during behavioral encoding. Indeed, the sustained firing during depolarizing current injections larger than 400 pA (this study) and in response to afferent input (Larimer and Strowbridge, 2010; Afrasiabi et al., 2022) are quintessential functional differences between GCs and SGCs. This sustained SGC firing is ideally suited to induce more immediate early gene (c-Fos or ARC) expression and could contribute to higher-than-expected labeling of SGCs during memory engram encoding. This is also supported by enrichment of activity dependent markers including PENK in behaviorally activated DG neurons, including SGCs labeled in TRAP2 mice (Erwin et al., 2020). Since the c-Fos-dependent ensemble labeling approach requires time for reporter expression, our experimental design does not allow a comparison of neuronal excitability at or before task performance. Nevertheless, our data demonstrating more sustained firing in SGCs and selectively reduced adaptation in labeled SGCs supports a role for greater neuronal activity in preferential recruitment of SGCs to task related dentate engrams.

It is possible that the sustained firing in SGCs, as well as higher NMDAR-mediated currents (Larimer and Strowbridge, 2010), contribute to their increased representation (∼30%) among behaviorally tagged DG neuronal ensembles compared to their relative population (∼3-5%) (Save et al., 2019). Furthermore, the wider dendritic arbors of SGCs are ideally positioned to receive distributed inputs and could support their preferential recruitment during behaviors. In this regard, whether SGCs and GCs differ in the inputs they receive from the entorhinal cortex is currently unknown. Although we find higher than expected SGCs among neuronal ensembles, the SGC labeling during BM and EE is considerably lower than the approximate 80% representation in DG engrams reported previously following novel environment exposure (Erwin et al., 2020). It is possible that restricting analysis to a subset of neurons filled during physiological recordings contributed to overrepresentation of SGCs among engram neurons in prior studies (Erwin et al., 2020). This is not surprising because SGCs, especially those in the sparsely populated molecular layer are more readily visualized and accessed for patch physiology than labeled GCs in the densely packed cell layer. Indeed, SGCs represent greater than 70% of the labeled neurons recorded in our study. Thus, our manual classification of sparsely labeled neurons by an expert investigator using previously validated morphometric features (Gupta et al., 2020; Afrasiabi et al., 2022) is likely to more accurately reflect the proportional labeling of SGCs.

Neurons in shared ensembles, including the DG to CA3 projections, show preferential connectivity and selective synaptic strengthening (Ryan et al., 2015; Rao-Ruiz et al., 2021). While connectivity between GCs is rarely observed in the healthy DG, whether SGCs with axon collaterals in the molecular layer make functional synaptic contacts on GC had not been examined. We leveraged findings that DG ensembles stably reactivate and engage downstream circuits up to 12 days after encoding (Kitamura et al., 2017) to evaluate local connectivity among SGCs and GCs in behaviorally recruited DG ensembles one week after encoding. Our paired recordings did not find evidence for glutamatergic connections between labeled SGCs and GCs, indicating that local DG engram refinement is not supported by mutual synaptic strengthening. Moreover, we observed no evidence of direct synaptic connectivity between GCs and SGCs regardless of whether the neurons were labeled or unlabeled. While it is possible that slice preparation could sever axon collaterals, we routinely record from cells over 50 µm below the surface and recovered extensive axon collaterals from SGC-GC pairs and excluded cells in which axon collaterals were not visualized. Moreover, SGCs have compact axonal distribution (Gupta et al., 2020), minimizing the possibility that lack of connections was a consequence of severed axons. Furthermore, even wide-field illumination to activate ChR2 positive terminals failed to evoke EPSCs in unlabeled GCs or SGCs, consistent with lack of connectivity between labeled and unlabeled SGC-GC pairs. Taken together with the evidence for increased overlap between DG ensembles labeled during encoding and recall, the limited glutamatergic interconnectivity among GCs and SGCs supports an instructive role for afferent inputs in DG ensemble recruitment. Indeed, we found that labeled neurons received more frequent and higher amplitude spontaneous glutamatergic inputs than corresponding unlabeled cells suggesting that strengthening of shared inputs may contribute to ensemble maintenance. Consistent with the proposal that afferent inputs contribute to DG ensemble recruitment, we found that the event timing of spontaneous glutamatergic inputs to pairs of labeled DG neurons were more correlated than inputs to labeled-unlabeled pairs suggesting that labeled neuronal pairs may receive correlated input streams. Moreover, input correlation was effective in discriminating between labeled-labeled versus labeled-unlabeled neuronal pairs, further supporting the role for input dependent recruitment of neuronal ensembles.

Lateral inhibition has been long considered a promising mechanism for discriminating amongst behaviorally relevant DG neuronal ensembles (Espinoza et al., 2018; Cayco-Gajic and Silver, 2019; Guzman et al., 2021; Borzello et al., 2023). Although GC mediated lateral inhibition of adjacent GCs is sparse (Espinoza et al., 2018; Braganza et al., 2020), focal activation of a random cohort of task-unrelated GCs, labeled with ChR2 by viral transfection, was shown to mediate surround inhibition of GCs (Stefanelli et al., 2016; Braganza et al., 2020). Moreover, SGCs, with their ability to robustly activate feedback inhibition (Larimer and Strowbridge, 2010; Afrasiabi et al., 2022), have been proposed as an ideal cell type to drive surround inhibition (Walker et al., 2010). However, unlike the findings based on focal activation of random GCs (Stefanelli et al., 2016), our analysis of SGCs and GCs tagged during naturalistic behavior found limited evidence for lateral inhibition of GCs. Even wide-field optical stimulation of labeled neurons, which would be expected to activate labeled terminals on interneurons due to the recruitment of intact and severed axons, rarely elicited lateral inhibition (1 of 55 trials). Our control experiments demonstrate that, unlike activation of sparse behaviorally labeled neurons, focal optical activation of cohorts of GCs labeled with ChR2 based on CAMKII expression supported robust feedback inhibition (Supplemental Fig. 3) in our slice preparation. Moreover, we have consistently recorded unitary IPSCs in DG interneuron-interneuron and interneuron-GC pairs (Yu et al., 2015; Yu et al., 2016; Proddutur et al., 2023) demonstrating that the circuit needed to support lateral inhibition is present in our slice preparation. Collectively, our results suggest that the sparse recruitment of behaviorally labeled ensembles may not be sufficient to elicit lateral inhibition. Our results are consistent with prior findings that focal activation of 2-4% of densely packed GCs is needed to recruit lateral inhibition in the DG (Braganza et al., 2020). Importantly, we identify that sparse behaviorally tagged ensembles are insufficient to support the kind of lateral inhibition observed during focal activation of high density virally labelled GCs (Stefanelli et al., 2016; Braganza et al., 2020 and Supplemental FIg. 3). Additionally, while modulating interneuron activity can constrain engram size by regulating network excitability (Morrison et al., 2016; Stefanelli et al., 2016), our microcircuit analyses suggest that SGCs have limited role in ensemble refinement by lateral inhibition. In this context, the possibility that slice recordings lead to underestimation of feedback dendritic inhibition cannot be ruled out. Curiously, we identified inhibition from a labeled GC to an unlabeled SGC and optically induced inhibition of an unlabeled GC, which does indicate presence of sparse lateral inhibition in the circuit. Overall, while there is considerable evidence for robust lateral inhibition in regulating DG activity, our data do not support the hypothesis that a sparse population of behaviorally active SGCs and GCs support ensemble refinement by surround inhibition.

The non-fear based contextual behavioral paradigms adopted in this study are known to engage the DG. However, they resulted in considerably sparser labeling than reported in contextual fear conditioning paradigms adopted in prior studies (Liu et al., 2012; Liu et al., 2014; Ryan et al., 2015; Roy et al., 2017). In order to specifically target neuronal cohorts activated during DG dependent spatial learning, we initially examined neuronal activation during the BM spatial navigation task involving spatial learning over multiple trials (Gawel et al., 2019). Tamoxifen treatment on day 6 labeled a sparse cohort of neurons, which was not compatible with circuit level analysis. Since the DG is preferentially activated by novelty (Hainmueller and Bartos, 2020; Mazurkiewicz et al., 2022; Borzello et al., 2023), we reasoned that learning related decrease in novelty may have contributed to sparse DG labeling during day 6 of BM spatial navigation. Consistent with a role for novelty in DG ensemble activation (Mazurkiewicz et al., 2022), tamoxifen induction during a single episode of enriched environment exposure reliably labeled a larger cohort of DG neurons, thereby enabling circuit analysis. Moreover, our demonstration of significantly greater co-labeling of tdT neurons with c-FOS upon a second exposure to the same environment, than following prior exposed to the BM, confirmed task specific neuronal labeling. While consistent with the rates of reactivation observed in fear conditioning experiments (DeNardo et al., 2019), c-FOS co-labeling was observed in less than 10% of the tdT positive neurons after re-exposure to the environment suggesting that not all tdT labeled neurons may be behaviorally relevant. A related caveat is the possibility that use of the TRAP2 system may miss active neurons expressing other IEGs (Heroux et al., 2018). Nevertheless, c-Fos driven labeling in TRAP2 mice remains the current best approach for activity dependent labeling, especially of DG neurons (Kawashima et al., 2014).

In summary, we find that SGCs represent about a third of dentate projection neurons labeled based on c-Fos expression during the contextual memory encoding, which is a considerable overrepresentation relative to their known population density. We propose that their unique sustained firing characteristics and temporal precision of afferent inputs may support their preferential labeling during activity dependent labeling of memory ensembles. Taken together, these data support a role for correlated inputs, the ability to sustain action potential firing, and sparse surround inhibition rather than glutamatergic interconnectivity as key determinants for recruitment of neurons to dentate memory ensembles.

## Materials and Methods

### Animals

All experiments were conducted under IACUC protocols approved by the University of California at Riverside and conformed with ARRIVE guidelines. c-Fos mice (TRAP2: Fostm2.1^(icre/ERT2)Luo/J^ ; Jackson Laboratories #030323) were back-crossed with C57BL6/N and were either bred with reporter line tdT-Ai14 mice (B6;129S6-Gt(ROSA)^26Sortm14(CAG-tdTomato)Hze/J^ ; Jackson Laboratories # 007908) to create TRAP2-tdT mice or reporter line Chr2-YFP (B6;129S-Gt(ROSA)^26Sortm32(CAG-COP4*H134R/EYFP)Hze/J^; Jackson Laboratories #12 569) to create TRAP2-ChR2/eYFP mice. Male and female TRAP2-tdT and TRAP2-ChR2/eYFP mice four to eight weeks old were used in experiments. Mice were housed with littermates (up to 5 mice per cage) in a 12/12 h light/dark cycle. Food and water were provided ad libitum.

### Behavioral training and engram labeling

Male and female experimental mice were trained in a spatial learning Barnes maze (BM) task or placed in an enriched environment (EE) for 3 hours followed by tamoxifen induction to induce Cre recombinase as detailed below. Since we observed TRAP2 mice exhibiting considerable litter to litter variability in tdT labeling following identical treatments in preliminary studies (not shown), we used littermate pairs for the following studies: ***Barnes maze (BM)***: 4-6 week old male and female TRAP2-tdT mice were trained in a spatial memory task on a Barnes maze table (Maze Engineers, https://conductscience.com/maze/), 92cm in diameter with 20 holes (5 cm diameter each). One hole was equipped with a false floor installed with a removable escape box that could be traded out for an additional false floor piece. The maze was set up in the middle of 4 curtain walls with two bright lights and a camera for recording above the maze. Different sets of visual cues (various shapes cut from felt) were attached to the curtain for spatial orientation. The escape hole was positioned in between two visual cues. Animals were held in their home cage outside of the curtain in a dark room until their turn to run the trial. We observed that these mice were hyperactive, therefore mice were housed individually on the day before training (Day 0). Mice were habituated to the behavior room in their home cages for at least 1 hour before training on Day 1, habituated to the arena by placing them in the starter cup on the table for 1 minute, and guided by gently moving the starter cup to the escape box (in a temporary location different from the experimental location). <u>During task acquisition training on</u> days 1-6, mice performed three 180-second trials during which the mouse explored the maze to find the escape box. The three trials were separated by a minimum of 15 minute inter-trial-intervals. If mice failed to locate the escape box at the end of the 180 seconds, the experimenter guided them to the escape box and then placed them back into their home cage. On Day 6 of BM acquisition, mice were brought to the room 5 hours before testing and received 4-hydroxy tamoxifen (4-OHT, 50mg/kg i.p.) 15 minutes prior to the first acquisition session. 4-OHT was prepared as described previously (DeNardo et al., 2019). Briefly, 4-OHT was dissolved in 100% ethanol at a concentration of 20mg/mL by sonicating solution at 37°C for 30 minutes or until dissolved, aliquoted and stored at −20°C. On the day of injection, 4-OHT was redissolved by sonicating solution at 37°C for 10 minutes. A 1:4 mixture of castor oil and sunflower seed oil, respectively, was added for a final concentration of 10mg/mL. The remaining ethanol in solution was evaporated by speed vacuuming in a centrifuge (DeNardo et al., 2019). Behavior in the Barnes maze paradigm was analyzed using Anymaze software by a blinded experimenter. Additionally, support vector machine-based, automated, Barnes Maze Unbiased Strategy (BUNS) classification algorithm and a non-arbitrary numerical cognitive score based on the BUNS analysis (Illouz et al., 2016) were used to evaluate the use of spatial strategy for Barnes Maze.

### Enriched environment (EE)

Experimental TRAP2-tdT and TRAP2-ChR2/eYFP mice were housed in an enriched environment consisting of an oversized cage filled with multiple tunnels, extra nestlets, a metal swing, and a few huts for the animals to interact with for 3 hours. Mice received 4-OHT (50mg/kg i.p) 90 minutes into their 3 hours of enrichment. Animals were left in the room for an additional 5 hours to limit neuronal activity labeling not related to the behavioral paradigm. In a subset of experiments (Fig. 1), 7 days following 4-OHT induction, littermate cohorts of TRAP2-tdT mice that underwent BM acquisition or EE exposure were placed with their respective pair into the EE for two hours and then immediately sacrificed by perfusion with 4% paraformaldehyde (PFA) upon removal from the EE. TRAP2-ChR2/eYFP mice induced after EE exposure were sacrificed a week later for electrophysiology (Fig. 2-5; Supplemental Fig. 2-4).

### Immunohistochemistry and Cell Morphology

TRAP2-tdT mice, 90 minutes following EE exposure, were transcardially perfused with PBS followed by a 4% PFA while under euthasol anesthesia. The brains were held in the 4% PFA at 4°C for 3 hours before being transferred to PBS. Coronal brain sections (50 μm) were obtained using a Leica vt100s vibratome and 5 sequential sections, each 250 μm apart across the septotemporal axis, were immuno-stained for c-Fos and analyzed for quantification. Free floating sections were blocked in 10% goat serum in PBS with 0.3% Triton X-100 for 1 hour. Sections were incubated in 4°C overnight in primary antibody for c-Fos (1:750, Rabbit mAb Cell Signaling Technology, cat #2250). The following day, sections were incubated in goat anti-rabbit Alexa Fluor® 488 secondary antibody (1:500 Abcam, cat #150077) for 1 hour.

Slices from TRAP2-ChR2/eYFP mice that were used in electrophysiological studies were fixed in 0.1mM phosphate buffer containing 4% PFA at 4°C overnight. Slices were washed with PBS and then incubated in 10% goat serum with 0.3% Triton X-100 for 1 hour at room temperature. Sections were incubated in 4°C overnight in primary antibody for GFP (1:500 Anti-Green Fluorescent Protein Antibody Aves Labs, AB_2307313). The following day, sections were incubated in goat anti-chicken Alexa Fluor® 488 secondary antibody (1:500 Abcam cat# 150169) and Alexa Fluor® 594 conjugated streptavidin (1:1000 Thermo Fisher, S11227) in PBS with 0.3% Triton X-100 for 2 hours at room temperature.

Slices were mounted on a glass slide using Vectashield®. Sections were imaged using a Zeiss Axioscope-5 with stereo investigator (MBF Bioscience) for analysis. Cell counts, cell type classification, and evaluation of double labeling were conducted by an experimenter blinded to treatments. Cells with compact dendritic arbors and somata with greater length than width were classified as GCs and those with wide dendritic angle, 2 or more primary dendrites, and greater somatic width than height were classified as SGCs (Gupta et al., 2020; Afrasiabi et al., 2022) by a trained investigator.

### Slice Physiology

Seven to nine days after tamoxifen induction following EE exposure, TRAP2-ChR2/eYFP mice were euthanized under isoflurane anesthesia for preparation of horizontal brain slices (350μm) using a Leica VT1200S Vibratome in ice cold sucrose artificial cerebrospinal fluid (sucrose-aCSF) containing (in mM): 85 NaCl, 75 sucrose, 24 NaHCO_3_, 25 glucose, 4 MgCl_2_, 2.5 KCl, 1.25 NaH_2_PO_4_, and 0.5 CaCl. Slices were bisected and incubated at 32°C for 30 min in a holding chamber containing an equal volume of sucrose-aCSF and recording aCSF and were subsequently held at room temperature for an additional 30 min before use. The recording aCSF contained (in mM): 126 NaCl, 2.5 KCl, 2 CaCl_2_, 2 MgCl_2_, 1.25 NaH_2_PO4, 26 NaHCO_3_, and 10 D-glucose. All solutions were saturated with 95% O_2_ and 5% CO_2_ and maintained at a pH of 7.4 for 2-6 hours (Gupta et al., 2012; Yu et al., 2015; Afrasiabi et al., 2022). Slices were transferred to a submerged recording chamber and perfused with oxygenated aCSF at 33°C. Whole-cell voltage-clamp and current-clamp recordings from GCs in the granule cell layer and presumed SGCs in the inner molecular layer or edge of the granule cell layer were performed under IR-DIC visualization with Nikon Eclipse FN-1 (Nikon Corporation) using 40x water immersion objective. Recordings were obtained using axon instruments MultiClamp 700B amplifier (Molecular Devices). Data were low pass filtered at 2kHz, digitized using Axon DigiData 1400A (Molecular Devices), and acquired using pClamp11 at 10kHz sampling frequency. Recordings were obtained using borosilicate glass microelectrodes (3-7MΩ), pulled using Narishige PC-10 puller (Narishige Japan). Recordings were performed using K-gluconate based internal solution (K-gluc) containing 126 mM K-gluconate, 4 mM KCl, 10 mM HEPES, 4 mM Mg-ATP, 0.3 mM Na-GTP, and10 mM PO-creatinine or cesium methane sulfonate (CsMeSO_4_) internal solution containing 140 mM cesium methane sulfonate, 10 mM HEPES, 5 mM NaCl, 0.2 mM EGTA, 2 mM Mg-ATP, and 0.2 mM Na-GTP. (pH 7.25; 270–290 mOsm). Biocytin (0.2%) was included in the internal solution for post hoc cell identification (Yu et al., 2015; Afrasiabi et al., 2022; Gupta et al., 2022). Cells labeled with eYFP were visualized under epifluorescence and patched under IR-DIC using pipettes filled with K-gluc internal and held at −70mV in current clamp. 10 ms, 10Hz pulses of blue light (λ=470 nm 0.9 mW) was used to optically evoke firing or inward currents were used to confirm ChR2/eYFP labeling. Responses to 1s positive and negative current injections, beginning at −200 pA with 40 pA steps up to 20 sweeps, were examined to determine active and passive characteristics. Dual patch clamp recordings were obtained from pairs of labeled and unlabeled neurons. Unlabeled neurons were recorded using microelectrodes with CsMeSO_4_ internal and held at 0mV (glutamate reversal potential) to isolate inhibitory postsynaptic currents (IPSCs) and −70mV (close to GABA reversal potential) to record excitatory postsynaptic currents (EPSCs). Labeled neurons, held in current clamp, were depolarized by 10ms 500pA pulses at 50Hz to elicit action potentials to measure evoked responses in labeled or unlabeled cells held in voltage clamp. Labeled neuron pairs were tested for connectivity in both directions. In control experiments, wild-type mice were bilaterally injected with AAV5-CaMKIIa-hChR2(H134A)-EYFP (gift from Karl Deisseroth, Addgene plasmid # 26969) in the granule cell layer (AP −3.2 mm, ML ±2.6 mm, DV −2.8 mm). Four weeks after injection, mice were sacrificed for horizontal hippocampal slices (350 µm thick) were prepared as detailed. Various regions of interest (ROIs) were selected for stimulation of ChR2 expressing cells GCs using a Digital Mirror Device (DMD)-based pattern illuminator (Mightex Polygon 400), coupled to 473 nm blue LED (AURA light source), and controlled via TTL based input from pClamp as detailed previously (Proddutur et al., 2023). Three progressively smaller circular ROIs with diameters of 110±10 µm, 55±5 µm and 27±3 µm were activated in the granule cell layer. Light intensity was set at 2.6 mW. Inhibitory postsynaptic currents (IPSCs) were recorded from granule cells outside the stimulated ROI using a cesium-based internal solution while holding the membrane potential at 0 mV.

Spontaneous EPSCs were (sEPSCs) were recorded in both labeled and unlabeled cells in slices from TRAP2-ChR2/eYFP and TRAP2-tdT mice induced with tamoxifen after EE and from C57BL6/N mice. Recordings were obtained from a holding potential of −70mV in voltage clamp for 5-10 minutes. In a subset of recordings, the sodium channel blocker tetrodotoxin (TTX 1 µM) was used to block action potential dependent events. In control experiments, sEPSC interevent interval (IEI) in aCSF was not different from the IEI recorded in GABA_A_ receptor antagonist, SR95531 (10 µM, IEI in aCSF in s: 5.29 ± 0.84, n=12 cells/3 mice; SR95531: 5.29 ± 0,98, n=9 cells/3 mice, p=0.7 by Mann Whitney U test) confirming that a majority of the synaptic events under these conditions are glutamatergic. Recordings were discontinued if series resistance increased by > 20% or if access resistance surpassed 25MΩ. Post hoc biocytin immunostaining and morphologic analysis was used to definitively identify SGCs and GCs included in this study.

### Data Analysis

Active and passive properties were analyzed using EasyElectrophysiology v2.6.3 (Easy Electrophysiology Ltd). Action potential, threshold, amplitude, halfwidth, and first spike latency were acquired from the first sweep in which the cell fired. Fast and medium afterhyperpolarization (AHP) and spike frequency adaptation were determined based on voltage response and firing in response to a 120 pA current injection. Action potential (AP) threshold was calculated using the first derivative method. Amplitude was calculated by the AP peak value minus baseline. Spike frequency accommodation was calculated using the divisor method, in which the inter spike interval (ISI) of the first two APs is divided by the ISI of the last two APs. First spike latency is the time from the start of a current pulse to the first AP.

Action potential and synaptic potential analysis were conducted using EasyElectrophysiology v2.6.3 (Easy Electrophysiology Ltd). Action potential kinetics were analyzed with a 200kHz interpolation for rise-time, decay-time, and halfwidth. Decay was measured using a biexponential decay curve fit with a cut off of 10-90% of the AP amplitude. Rise time is calculated between 10-90% of the AP amplitude. Half width was calculated as the time between the two half-amplitude samples. Afterhyperpolarization values are calculated as baseline minus fAHP or mAHP. The value is the minimum point within a search region specified as 0-3ms for fAHP and 10-50ms for mAHP. Spontaneous EPSCs were detected and analyzed using EasyElectrophysiology threshold search algorithm and events were confirmed by the experimenter. Any “noise” that spuriously met trigger specifications was rejected. Cumulative probability plots in Fig. 5 were obtained using the same number of events from each cell.

Temporal Correlation of sEPSCs: Synaptic event times used for temporal correlation analysis were extracted from sEPSCs recorded at a holding potential of −70 mV in low chloride internal solution. Temporal correlation of sEPSCs in dual recording sessions from cell pairs (labeled to labeled (L-L) and labeled to unlabeled (L-U)) was defined by a session-wise cross correlation profile (CCP) of temporally binned data for select detection windows (MATLAB *xcorr*, Wiegand & Cowen). Temporal correlations of sEPSC event times in a large ±1 s detection window confirmed an expected high cross correlation of events within the 100 ms central bin. A small ±10 ms detection window with 1 ms bin width resulted in too few cooccurring events. Consequently, cross correlation profiles (CCPs) were developed using multiple *detection windows* (±10ms to ±1s) with corresponding *bin durations* (21 bins within window). Temporal correlation across full session timelines was not calculated to avoid spuriously high correlation values from simultaneous absence of events in cell pairs (Cutts and Eglen, 2014). As an additional measure to avoid specious correlations, sessions with too few events, low event frequency, and short recording durations were not analyzed. Temporal correlations were tested using detection windows and bin sizes were always divided into 21 equally sized bins in the window: a ±100 ms detection window with 10 ms bins (200 ms window, 21 bins aligned to sEPSC) and a ±50 ms detection window with 5 ms bins (100 ms window, 21 bins aligned to sEPSC). The dynamics of the cross-correlograms generated from our data sets using previously established methods to evaluate monosynaptic connectivity (Bartho et al., 2004; Senzai and Buzsaki, 2017) parallelled that of the CCP plots (Supplemental Fig. 6) illustrating that the methods similarly capture co-dependencies between event time series.

Temporal correlation was determined if the CCP exceeded a 2 standard deviation (SD) threshold above the total mean correlation (0.15 for the 100 ms detection window and 0.10 for the 50 ms detection window). Within each detection window, “peri-occurrence” was defined by the CCP maximum outside the center bin while “cooccurrence” was defined by the correlation in the center bin. CCP of sEPSC event times in L-L and L-U pairs were compared with randomly jittered event times from the same data set to identify intrinsic correlations within the data. Session jitter was pseudo-randomly selected from an event timeline matrix (assigned to one cell from each paired cell recording session) bound by ± 0.5 s across 100 iterations. The temporally jittered correlation data were then compared to the temporally aligned CCP and correlation data in the center bin (Wiegand et al., 2016). The ability of the sEPSC event cooccurrence to predict L-L versus L-U pairs was computed by plotting the receiver operating characteristic (ROC) curve and calculating the area under the curve (AUC) in both groups. Correlation values in the center bin of the CCPs were used to generate histograms (correlation bins from 0 to 0.5 with correlation bin widths of 0.001), which were reverse integrated to evaluate the cumulative sum between categorized L-L sessions (true, n=7) and L-U sessions (false, n=8) rates. Cumulative sums were used to find the total AUROC as the classification performance measure using MATLAB (*hist counts*, *cumsum*, *flip*, and *trapz* functions), where 50% AUROC performance would classify L-L versus L-U by random chance.

Sample sizes were not predetermined and conformed with those employed in the field. Significance was set to p<0.05, subject to appropriate Bonferroni correction. Statistical analysis was performed using GraphPad Prism 10 and MATLAB. Data were tested for normality and unpaired Kolmogorov-Smirnov (KS) Test, unpaired Mann-Whitney, one-way ANOVA, two-way ANOVA (TW-ANOVA), two-way repeated measures ANOVA (TW-RM-ANOVA), or Kruskal-Wallis followed by post-hoc pairwise multiple comparisons using Holm-Sidak method or Dunn’s method used as appropriate. Statistical data is reported as mean ± SEM or median (interquartile range) as appropriate.

## Acknowledgements

This work is supported by National Institutes of Health (NIH) NINDS R37NS069861, R01NS097750 to V.S., NIH/NINDS F31NS124290 to L.D.

## Author contributions

L.D, M.A performed experiments; L.D, M.A, K.M and E.Z analyzed data; L.D, M. A, K.M. E.Z and V.S. interpreted results of experiments; L.D, M.A and K.M. prepared figures; L.D, and V.S. conception and design of research; L.D., K.M. and V.S. drafted manuscript; L.D., M.A, K.M. E.Z and V.S. finalized and approved manuscript.

