## Supplementary Figure and Table for "Cellular and circuit features distinguish dentate gyrus semilunar granule cells and granule cells activated during contextual memory formation"

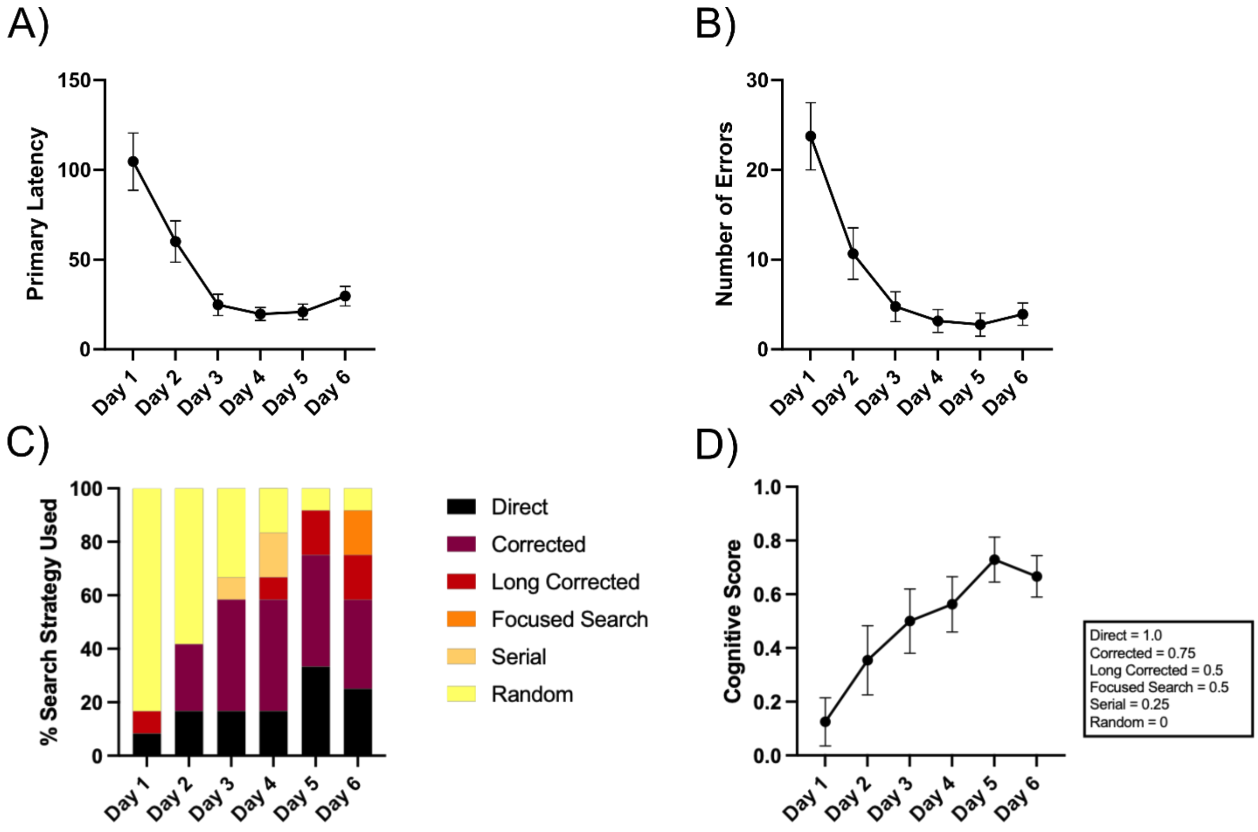


***Supplemental Figure 1: Search strategies adopted in the Barnes maze task.*** *A-B) Plot of primary latency (A) and number of errors (B) to find the escape hole over training days. C)Visualization of search strategies used in the Barnes maze paradigm. D) Summary cognitive score based on search strategy.*

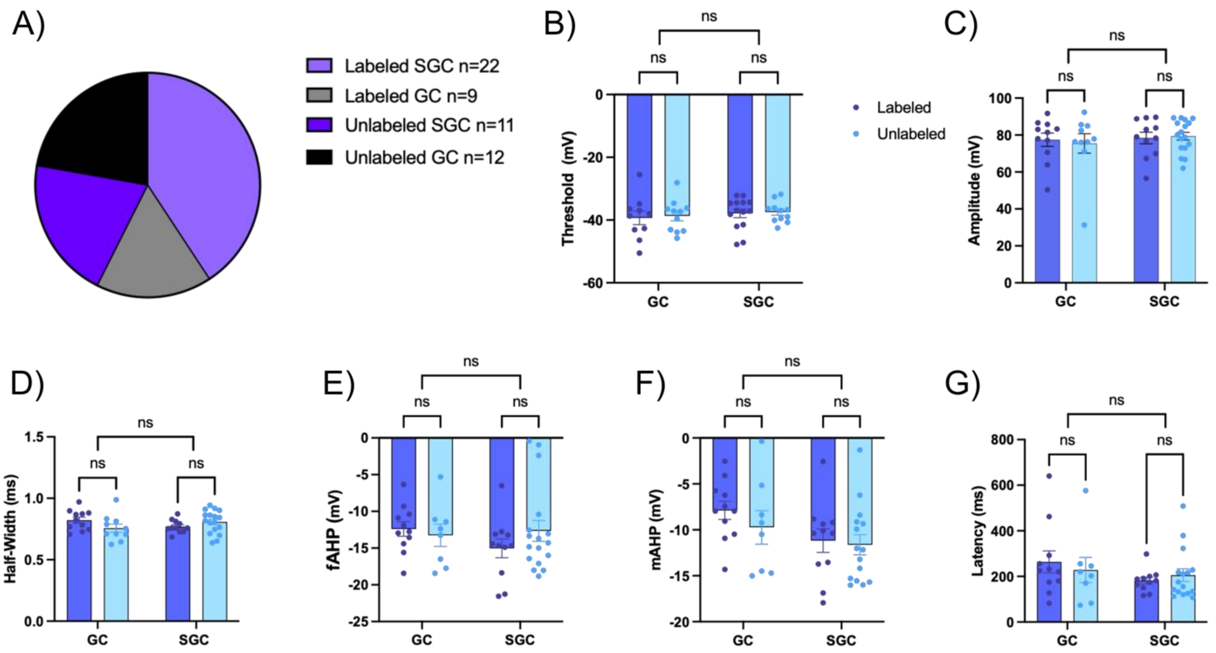


***Supplemental Figure 2: Active properties of labeled and unlabeled GCs and SGCs.*** *A) Pie chart showing the proportion of labeled and unlabeled GCs and SGCs included for analysis of active membrane properties. Note the greater proportion of SGCs represented among labeled neurons. B-G) Summary histograms of threshold of action potential (B), amplitude (C), half-width (D), fast (E) and medium afterhyperpolarizations (F) and latency (G). Data are presented as mean ± SEM.*

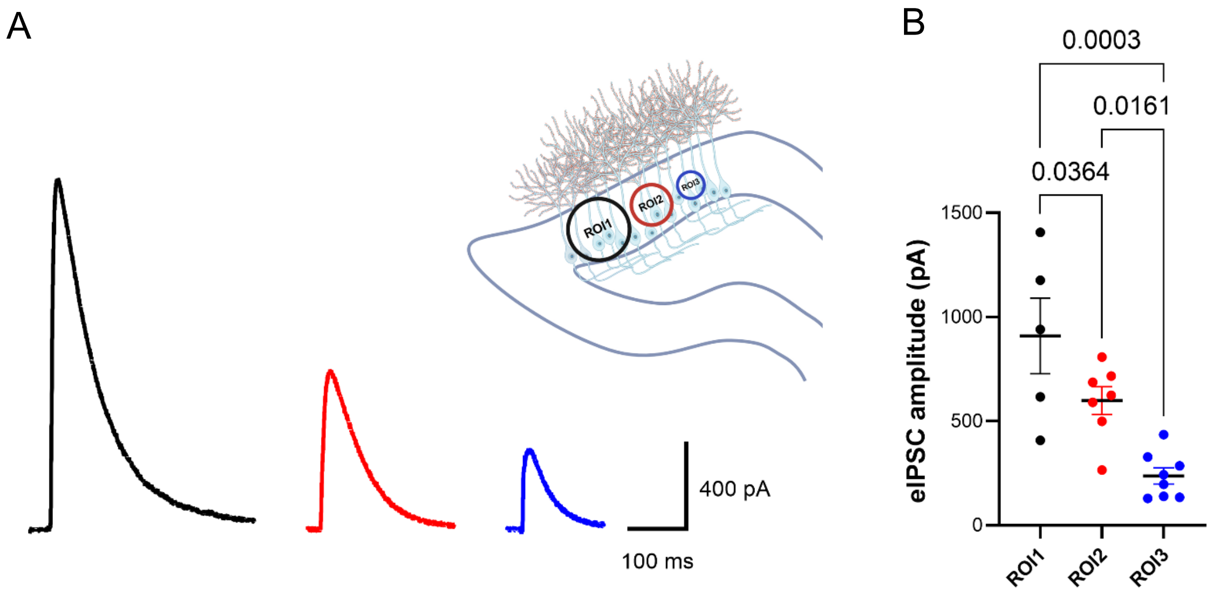


***Supplemental Figure 3. Robust feedback inhibition in response to focal activation of a random cohort of granule cells.*** *A). Example optically evoked IPSCs in slices from mice injected with* AAV5-CaMKIIa-hChR2(H134A)-EYFP *in response to activation of three progressively smaller regions of interest (ROIs), the largest spanning the granule cell layer. Inset: Schematic of ROI selection in the DG . B) Summary plot of eIPSC amplitude in response to optical activations of the three ROIs. P values indicated in the one-way ANOVA, with a significance threshold set at $p < 0.05$. Recordings were obtained from 5-8 cells from 3 mice.*

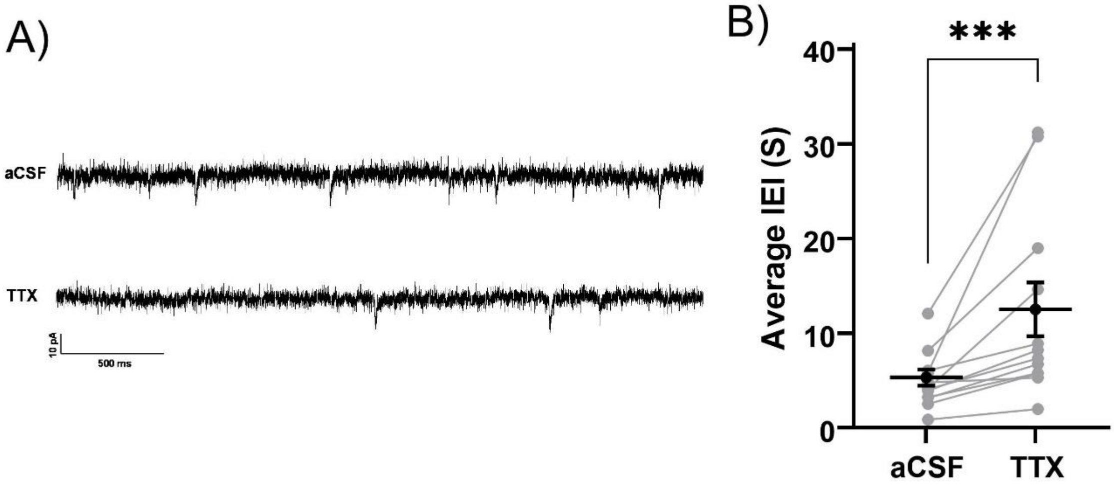


***Supplemental Figure 4: Spontaneous EPSCs in dentate GCs include action potential driven events.*** *A) Representative current traces from a GC illustrates spontaneous EPSCs (in aCSF, above) and miniature EPSCs (in TTX, below). B) Summary of EPSC interevent interval (IEI) in aCSF and after perfusion of TTX. Data presented as mean ± SEM. *** indicates p=0.0005 by paired t-test.*

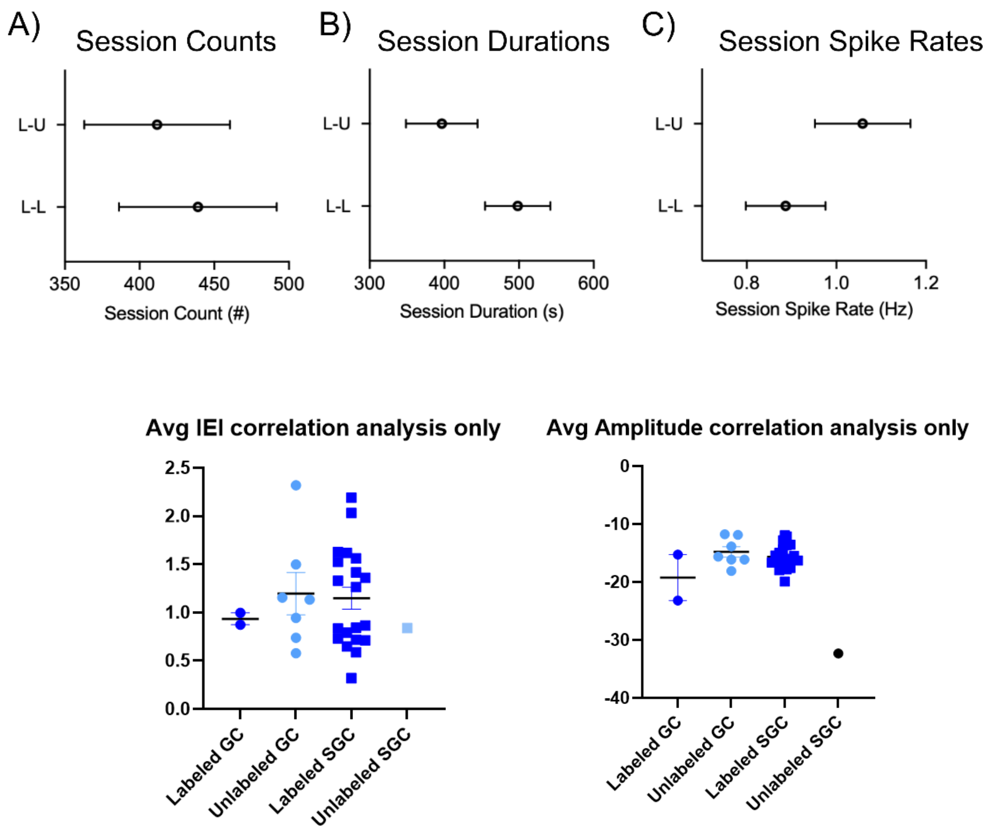


***Supplemental Figure 5: L-L and L-U Sessions do not differ in event rates.*** *A) sEPSC event counts (LL n=14, LU n=16), B) Recording durations(LL n=7, LU n=8) and C) event frequency (LL n=14, LU n=16) for data used in correlation analysis. Data presented as mean ± SEM. D-E) Distribution of average sEPSC inter-event interval (D) and amplitude (C) in labeled and unlabeled GCs and SGCs included among the LL and LU pairs.*

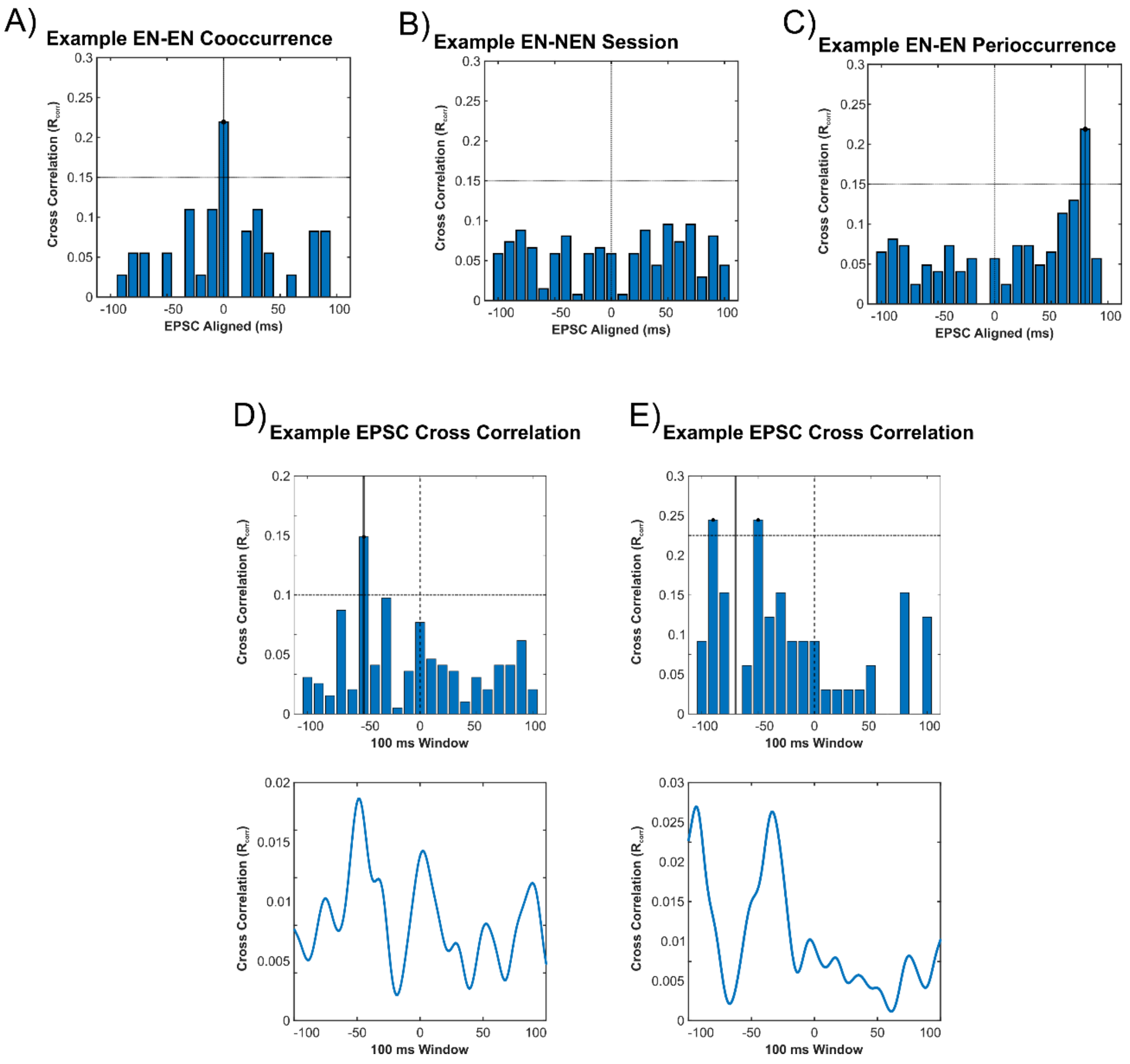


***Supplemental Figure 6*****: *Example sEPSC cross correlation profiles.*** *A) Representative CCP for L-L dual recording session with maximum correlation in the center bin (cooccurrence). B) Representative CCP for L-U dual recording session with no maximum correlation within detection window (no coincidence, same session as in Fig 5C). C) Representative cross correlation profile (CCP) of sEPSCs for L-L dual recording session with maximum correlation within detection window (peri-occurrence, same session as in Fig 5B). D-E) xamples of EPSC cross-correlation histograms generated using the CCP method (above) and using the MATLAB cross-correlation function xcorr (below).*

### *Dovek et al Supplemental Table*

*Supplemental Table 1 Electrophysiological properties of labeled and unlabeled, SGCs and GCs.*

| Parameter (Unit) | Labeled GC (GC-L) | Unlabeled GC (GC-U) | Labeled SGC (SGC-L) | Unlabeled SGC (SGC-U) | F & P value | Multiple Comparisons |
|---|---|---|---|---|---|---|
| Threshold (mV) | -39.31 ±2.181 | -38.69±1.552 | -37.87 ±1.342 | -37.47±0.9952 | Interaction: F (1, 42) = 0.004967 P=0.9441 | GC-L vs GC-U p=0.9978 |
| | | | | | Cell Type: F (1, 42) = 0.7407 P=0.3943 | SGC-L vs SGC-U p=0.9994 |
| | | | | | Labeling: F (1, 42) = 0.1120 P=0.7395 | GC-L vs SGC-L p=0.9412 |
| | | | | | by Two way ANOVA | GC-U vs SGC-U p=0.9703 |
| Amplitude (mV) | 75.44±5.275 | 77.44±3.570 | 79.37±2.109 | 78.43±3.137 | Interaction: F (1, 45) = 0.1842 P=0.6698 | GC-L vs GC-U p=0.327 |
| | | | | | Cell Type: F (1, 45) = 0.5138 P=0.4772 | SGC-L vs SGC-U p=0.7098 |
| | | | | | Labeling: F (1, 45) = 0.02396 P=0.8777 | GC-L vs SGC-L p=0.4775 |
| | | | | | by Two way ANOVA | GC-U vs SGC-U p=0.5133 |
| Half-Width (ms) | 0.7572±0.03330 | 0.8219±0.02489 | 0.8076±0.02261 | 0.7700±0.01597 | Interaction: F (1, 45) = 4.130 P=0.0481 | GC-L vs GC-U p=0.3227 |
| | | | | | Cell Type: F (1, 45) = 0.0009773 P=0.9752 | SGC-L vs SGC-U p=0.7098 |
| | | | | | Labeling: F (1, 45) = 0.2903 P=0.5927 | GC-L vs SGC-L p=0.4775 |
| | | | | | by Two way ANOVA | GC-U vs SGC-U p=0.5133 |
| fAHP (mV) | -13.26±1.519 | -12.41±0.9795 | -12.66±1.426 | -15.04±1.272 | Interaction: F (1, 43) = 1.269 P=0.2662 | GC-L vs GC-U p=0.9919 |
| | | | | | Cell Type: F (1, 43) = 0.5024 P=0.4823 | SGC-L vs SGC-U p=0.5947 |
| | | | | | Labeling: F (1, 43) = 0.2822 P=0.5980 | GC-L vs SGC-L p=0.9972 |
| | | | | | by Two way ANOVA | GC-U vs SGC-U p=0.5905 |
| mAHP (mV) | -9.731±1.826 | -7.885±0.9965 | -10.06 ±1.447 | -11.17 ±1.279 | Interaction: F (1, 41) = 0.2897 P=0.5933 | GC-L vs GC-U p=0.8233 |
| | | | | | Cell Type: F (1, 41) = 4.049 P=0.0508 | SGC-L vs SGC-U p=0.9979 |
| | | | | | Labeling: F (1, 41) = 0.7999 P=0.3763 | GC-L vs SGC-L p=0.7728 |
| | | | | | by Two way ANOVA | GC-U vs SGC-U p=0.2675 |
| Latency (ms) | 228.2±55.35 | 262.9±47.98 | 205.5±27.86 | 181.2±14.79 | Interaction:F (1, 42) = 0.6229 P=0.4344 | GC-L vs GC-U p=0.9579 |
| | | | | | Cell Type: F (1, 42) = 1.955 P=0.1694 | SGC-L vs SGC-U p=0.9784 |
| | | | | | Labeling: F (1, 42) = 0.01930 P=0.8902 | GC-L vs SGC-L p=0.9883 |
| | | | | | by Two way ANOVA | GC-U vs SGC-U p=0.4179 |
| Spike Frequency Accommodation | 0.3271±0.07460 | 0.2791±0.05569 | 0.7846±0.07658 | 0.45657±0.06271 | Interaction: F (1, 41) = 3.477 P=0.0694 | GC-L vs GC-U p=0.9876 |
| | | | | | Cell Type: F (1, 41) = 19.66 P<0.0001 | SGC-L vs SGC-U p=0.0064 |
| | | | | | Labeling: F (1, 41) = 6.379 P=0.0155 | GC-L vs SGC-L p=0.2611 |
| | | | | | by Two way ANOVA | GC-U vs SGC-U p=0.0003 |
| Input Resistance (MOhms) | 143.8 ± 8.673 | 169±13.51 | 115±5.974 | 128.4±7.817 | Interaction: F (1, 48) = 0.4385 P=0.5110 | GC-L vs GC-U p=0.2365 |
| | | | | | Cell Type: F (1, 48) = 14.98 P=0.0003 | SGC-L vs SGC-U p=0.7091 |
| | | | | | Labeling: F (1, 48) = 4.660 P=0.0359 | GC-L vs SGC-L p=0.0760 |
| | | | | | by Two way ANOVA | GC-U vs SGC-U p=0.0155 |
